# Flexible 3D *kirigami* probes for *in vitro* and *in vivo* neural applications

**DOI:** 10.1101/2024.11.05.622167

**Authors:** M. Jung, J. Abu Shihada, S. Decke, L. Koschinski, P. S. Graff, S. Maruri Pazmino, A. Höllig, H. Koch, S. Musall, A. Offenhäusser, V. Rincón Montes

## Abstract

Three-dimensional (3D) microelectrode arrays (MEAs) are gaining popularity as brain-machine interfaces and platforms for studying electrophysiological activity. Interactions with neural tissue depend on the electrochemical, mechanical, and spatial features of the recording platform. While planar or protruding two-dimensional MEAs are limited in their ability to capture neural activity across layers, existing 3D platforms still require advancements in manufacturing scalability, spatial resolution, and tissue integration. In this work, we present a customizable, scalable, and straightforward approach to fabricate flexible 3D *kirigami* MEAs containing both surface and penetrating electrodes, designed to interact with the 3D space of neural tissue. These novel probes feature up to 512 electrodes distributed across 128 shanks in a single flexible device, with shank heights reaching up to 1 mm. We successfully deployed our 3D *kirigami* MEAs in several neural applications, both *in vitro* and *in vivo*, and identified spatially dependent electrophysiological activity patterns. Flexible 3D *kirigami* MEAs are therefore a powerful tool for large-scale electrical sampling of complex neural tissues while improving tissue integration and offering enhanced capabilities for analyzing neural disorders and disease models where high spatial resolution is required.

## 1. Introduction

Unveiling the intricate networks of the central nervous system and their functions in both healthy and pathological states requires investigating neural activity with high spatiotemporal resolution across multiple topological levels. This includes examining activity from individual neurons to local and regional dynamics within distinct neural structures and encompasses the three-dimensional (3D) space of nervous tissues, extending from its surface to its complex, multilayered internal structure. Neuroelectronic interfaces comprising microelectrode arrays (MEAs) facilitate this by capturing electrical signals with sub-milliseconds resolution and covering spatial regions from micrometers to millimeters. MEAs enable the recording of complementary neural information by capturing high-frequency signals associated with action potentials (also known as spiking activity), and low-frequency signals related to local field potentials (LFPs), which represent the activity of individual neurons and the collective activity of neuronal populations in localized neural areas, respectively. Consequently, MEAs are commonly used to record extracellularly and modulate the electrical activity of various neural structures in basic neuroscience and pre-clinical and clinical settings. Neuroscientific applications of MEAs range from understanding the neural basis of memory^1^, space^2^, and behavior^3^ to studying neural oscillations in epilepsy research^4^ and developing brain-machine interfaces (BMIs) to restore lost sensorimotor functions^5^.

MEAs containing grids of microelectrodes spatially arranged in two dimensions on stiff or flexible flat surfaces (2D MEAs) are widely used to capture neural dynamics from the surface of nervous tissues. 2D MEAs enable the study of neural slices *in vitro*, such as the brain^6^ and retina explants^7^, as well as *in vivo* applications, such as capturing neural oscillations with electrocorticography arrays on the brain surface^8^ or interfacing with the spinal cord using paddle electrodes^9^. To access the deeper intraneural space, penetrating MEAs are employed. These include needle-like arrays, such as the Utah array^10^ or microwire arrays^11^, and comb-like arrays with multiple shanks, such as the Michigan array^12^ or thin-film polymer-based shank (also referred to as thread or strip) arrays^13^. Furthermore, multi-contact single shank probes are also commonly used, including depth electrodes for deep brain stimulation or stereoelectroencephalography, as well as high-density probes based on complementary metal oxide semiconductor (CMOS) technology, such as Neuropixels probes^14^. While needle-like arrays provide access to the internal structures of nervous tissues, they typically capture information from a single neural layer across a millimeters-wide region. Although the electrode sites protrude, they are spaced at the same depth, resulting in only a two-dimensional (2D) coverage of the target region. Similarly, comb-like MEAs and single-shank devices enable either a 2D or a one-dimensional (1D) sampling of the tissue, with shanks that cover multiple layers at one or a couple of spatial locations (depending on the number of shanks).

To interact with 3D neural space, various types of 3D MEAs have been developed to penetrate and reach different neural layers and regions by implementing variations or extensions of 1D or 2D MEA technologies. Based on silicon (Si) technology, Utah and Michigan arrays have long been the gold standard for BMIs. Innovations such as the Utah slant array, which features needles at varying depths^15,16^, and the stacking and overlaying of 2D Michigan arrays with predefined spacers^17–20^, have enabled 3D spatial sampling of nervous tissues. However, these devices often trigger strong foreign body reactions (FBRs) due to the mechanical mismatch between the stiff Si and the soft neural tissue^21,22^.

Significant progress has been made in recent years with the development of softer, more flexible materials and micro-scale electrodes that can be used in 3D configurations while minimizing FBRs^23,24^. These devices typically involve the individual implantation of multiple flexible, penetrating threads or shanks, allowing coverage of different neural layers and regions, thereby enabling 3D sampling of the intraneural space^25–27^. However, this method does not allow simultaneous access to both surface and intraneural activity, making it time-consuming since each thread must be individually addressed, significantly increasing implantation time or requiring the use of sophisticated robots for automated implantation. Additionally, recreating the relative positions of individual 1D or 2D probes further increases the complexity of subsequent data analysis, especially because individual probes are prone to displacement due to micromotions, such as respiratory or cardiovascular movements. Moreover, flexible needle-like probes with varying heights and multisite designs per needle have been demonstrated^28,29^; But increasing electrode density of individual needles is limited by the resolution and scalability of additive manufacturing processes typically proposed in these works. Furthermore, simultaneous access to surface and deep neural activity has been demonstrated in experimental settings by utilizing surface and penetrating MEAs concurrently. This method strongly increases the handling and experimental complexity, as it requires the insertion of two independent devices in the same anatomical structure^30^.

A potential approach to overcome these existing challenges is the development of multi-contact and fully flexible 3D MEAs. Several research groups have proposed *kirigami*-based methodologies to achieve this goal. Inspired by the Japanese art of cutting and folding paper, *kirigami* techniques are adapted to micromachining by first processing a 2D flexible MEA. The 2D MEA is then etched to create cut-out patterns, allowing the shanks to be lifted into a 3D configuration. Various folding methods have been explored, including the use of ferromagnetic sheaths to induce folding *via* magnetic fields^31^, as well as electrostatic^32^ or mechanical actuation^33,34^ for selective shank folding.

While these existing 3D flexible *kirigami*-based MEAs have successfully enabled 3D neural sampling, from *in vitro* 3D neuronal cultures^33^ to simultaneous cortical and intracortical monitoring *in vivo*^34^, current fabrication techniques face scalability limitations. Manual shank lifting is time-consuming, and the use of ferromagnetic materials can involve cytotoxic metals such as nickel, along with the need for strong magnetic fields (*e.g.*, 100–400 mT) that may pose safety risks^31^. Additionally, these methods can lead to inconsistent folding angles or shanks folding back after implantation, limiting the precision and reliability of the resulting 3D structure^33^.

To overcome the spatial sampling and manufacturing limitations of 3D flexible MEAs, we developed a flexible, biocompatible, and high-resolution *kirigami* MEA using a parallelized folding technique based on a matched die forming process. This approach enables rapid, scalable, and customizable fabrication of 3D flexible MEAs with up to 128 shanks, providing a straightforward and reproducible method for producing complex neural interfaces. We demonstrate the versatility of our technique by fabricating 3D *kirigami* MEAs with various designs tailored for diverse neural applications, both *in vitro* and *in vivo*. Furthermore, we showcase the utility of these devices in neuroscientific studies, including the observation of seizure-like activity in human brain slices from epileptic patients and measuring sensory responses in the somatosensory and visual cortex in live mice. Together, this work demonstrates that our *kirigami*-based method provides a powerful new tool for capturing the multiregional and multilayered complexity of diverse neural tissues.

## 2. Results and Discussion

### 2.1. Fabrication of 3D flexible Kirigami neural probes

We developed a process to reliably fabricate 3D flexible MEAs based on the k*irigami* principle of cutting and folding a 2D design into a 3D structure using flexible polymeric materials, such as parylene-C (PaC). The process consists of three main steps: fabrication of a 2D *kirigami* template (Figure 1A), folding of the 2D *kirigami* template into its 3D conformation (Figure 1B), and thermoforming of the 3D flexible structure (Figure 1C). To fabricate the 2D *kirigami* template, we employed a mechanically robust 2D *kirigami* design and utilized standard surface micromachining processes and thin-film technology. We then used an additive manufacturing process to fabricate a customized mold containing upright structures that matched the location of the penetrating shanks in the 2D design. The mold enabled a matched die forming system to align and fold the 2D *kirigami* template in the third dimension. Lastly, the device was thermoformed to maintain the newly formed 3D structure. This process stands out as it allows scalability towards the parallel folding of multiple shanks (up to 512 tested so far) with a yield of 98%, avoiding the use of toxic materials or hazardous processes as proposed by other works^31,32^. Likewise, the process offers design customizability and flexibility, making it possible to adapt designs across a diverse range of spatial resolutions and neuronal models for both *in vitro* and *in vivo* applications.

**Figure 1:**
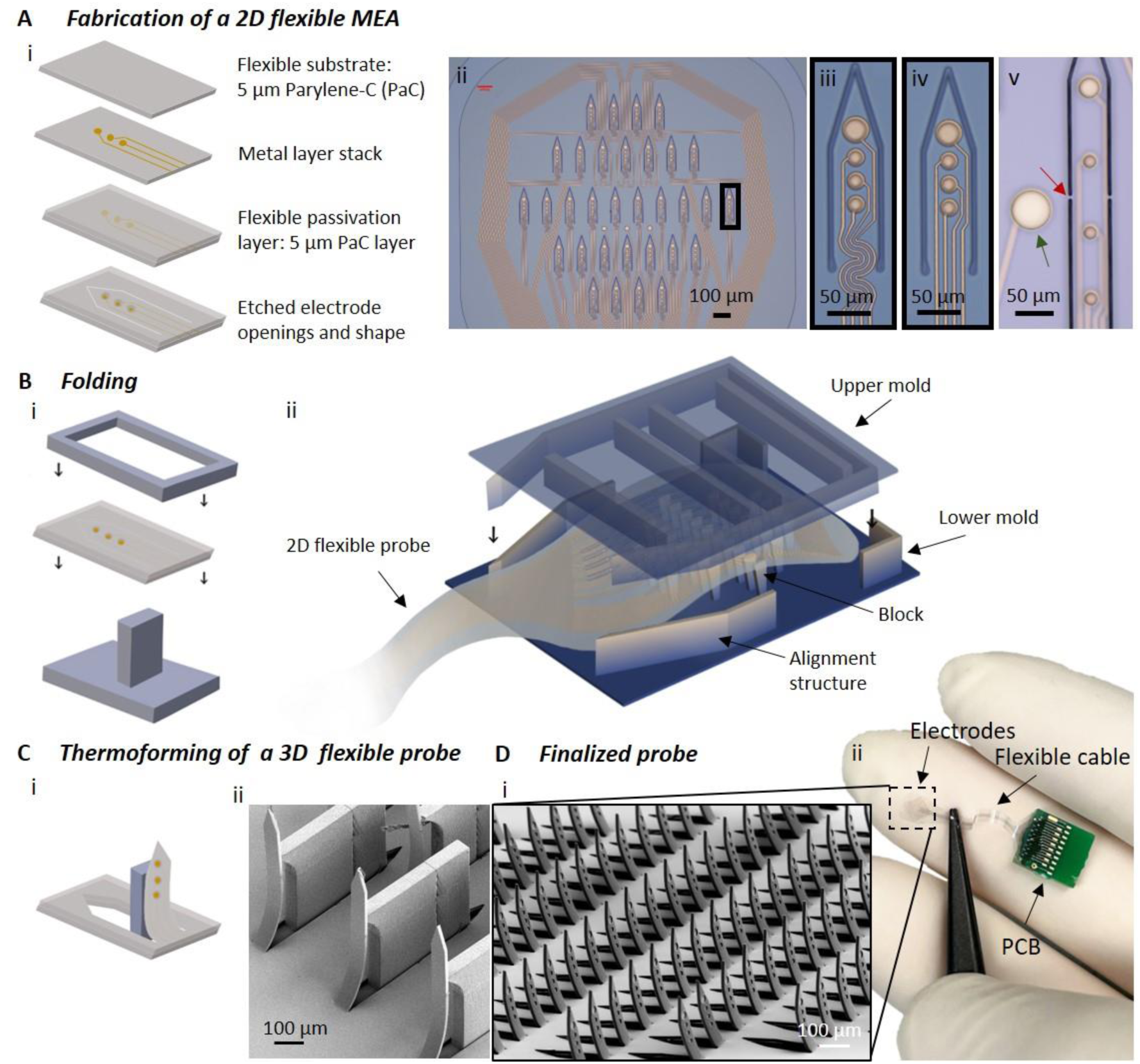
Fabrication of 3D *kirigami* MEAs. A) The fabrication of the 2D *kirigami* template comprises four steps (A_i_): 1) Chemical vapor deposition (CVD) of a 5 µm PaC layer (in grey). 2) Deposition of a Ti/Au/Ti metal layer stack (in yellow). 3) 2^nd^ CVD process for a PaC passivation layer. 4) Reactive ion etching (RIE) to open the electrodes and etch the shape. Exemplary design aimed at retinal applications with 32 shanks (A_ii_) containing meander-shaped (A_iii_) or straight feedlines (A_iv_). Additionally, surface electrodes (A_v_, green arrow) and breaking points for long and/or thin shanks (A_v_, red arrow) are included in the design. B) For the folding of the shanks, the 2D *kirigami* template is placed between a lower and an upper mold (one shank displayed in (B_i_) and complete probe in (B_ii_)) folding all shanks at the same time by blocks which are arranged at each shank location and creating homogenous pressure over the complete flexible template using the upper mold. C) The flexible probe placed inside the lower mold (one shank in (C_i_) and scanning electron microscopy (SEM) picture of several shanks in (C_ii_)) is then thermoformed. D) The finalized probe (SEM picture in (D_i_)) bonded on a PCB in comparison to a human hand (ii).

#### 2D kirigami template

The process starts with the fabrication of a 2D *kirigami* template, which features a flexible sandwich structure that ultimately forms a flexible 2D MEA. This structure is composed of two PaC layers that serve as substrate and encapsulation, and a metal stack layer of Ti/Au/Ti in the middle that shapes the electrical feedlines and electrodes (Figure 1A_i_). The design of the overall probe included a contact pad region, a flexible cable, and a sensing area containing surface electrodes and penetrating shanks, each holding multiple microelectrodes (Figure S1). Besides the standard electrode and contact pad openings and the outline of the implant, the 2D pattern contained the cut-outs corresponding to the structures to be folded upright in the third dimension (Figure 1A_i__i-iv_). The 2D design was then patterned *via* photolithography and further etched *via* dry etching, resembling the drawing and cutting steps of the *kirigami* process.

For a successful parallelized *kirigami* approach, the design of the 2D pattern served as a template for the 3D structure to be formed and also provided mechanical stability during the folding process. To reduce mechanical stresses at the folding sites of each shank, meander-shaped feedlines were integrated into the design at the kink (Figure 1A_iii_) as an alternative to straight feedlines (Figure 1A_iv_), and breaking points were added at the sides of each shank to provide mechanical stability to the 2D template (Figure 1A_v_, red arrow). The latter design strategy was critical for the success of the matched die forming process with long shanks above 300 µm and thin film layers below 10 µm, as it prevents high-aspect ratio and thin structures from failing due to self-bending or moving out of place due to electrostatic forces. Additionally, to increase the spatial resolution of the implant, electrodes were added on the plane surface of the 2D *kirigami* template next to or between the shanks.

### Folding of the 2D kirigami template

After microfabrication, the 2D *kirigami* template, also referred to as 2D flexible MEA, was flip-chip bonded onto a printed circuit board (PCB). The device was then ready for folding. Here, customized molds that matched the design of the 2D *kirigami* template were fabricated by two-photon lithography. Using complementary dies, hereafter referred to as upper and lower molds, we implemented a matched die forming process (Figure 1B and S2). The lower mold contained protruding blocks that exactly matched the location and geometry of the shanks in the 2D *kirigami* template and sidewalls performing as alignment structures that matched the outline of the sensing area of the 2D flexible MEA (Figure 1B_i_ and 1B_ii_). For short shanks, the block height surpassed the shank length, resulting in block height-to-shank length ratio of 1.11 (*e.g.*, 250 µm block height for 225 µm shank length). In contrast, for longer shanks, the blocks were shorter than the shank length, leading to ratio below 1 (*e. g.* 750 µm block height for 1000 µm shank length). To allow tolerance for the removal of the mold in further steps, the width of the blocks was 0.9-0.92 times the width of the shanks. Additionally, ramped side-walled blocks were implemented to reduce the contact surface area between the blocks and the shanks, facilitating the releasing step (*e.g.*, 120 µm at the top, 60 µm at the bottom).

This mold design allows the straightforward placement of the 2D *kirigami* template into the lower mold without requiring a complex alignment setup. Complementary to the lower mold and the 2D *kirigami* template, the upper mold contained straight protruding structures perpendicular to the planar axis of the shanks and located between rows of shanks and along the outer boundaries, thereby ensuring homogenous pressure on the 2D *kirigami* template when pressed towards the lower mold. The outer boundaries of the upper mold fit perfectly inside the lower mold (Figure 1Bii, tolerance to each side 35 µm) which ensures facile alignment, and together with the matching design of the lower mold, a matched die compression is enabled, allowing the concurrent folding of all shanks (Supplementary Video SV1).

### Thermoforming of a 3D kirigami MEA

Given that PaC, a thermoplastic, was used as the main substrate and encapsulation material, the flexible probe was subjected to a thermoforming step while still placed inside the lower mold after the folding (Figure 1C_i_ and 1C_ii_). Thermoplastics are moldable materials when subjected to temperatures above their glass transition temperature (*e.g.*, 90°C for PaC) and below their melting point (*e.g.*, 290°C for PaC)^35^. At high temperatures, the chain structures of the amorphous regions of the PaC undergo rearrangement, softening the material and enabling the reshaping of its molecular structure. The polymeric material then retains the new shape upon cooling and removal of the mold. The thermoforming protocol was optimized considering the trade-off between temperature and thermoforming time. While higher temperatures allowed proper thermoforming, they caused the material to stick to the molds. On the other hand, the use of lower temperatures implied longer thermoforming periods that in most cases were not fully met, causing then the shanks to fold back immediately after the separation from the mold. In the case of PaC, it had to be considered that high temperatures (> 300 °C) permanently change the mechanical and optical properties of the material^36^. Hence, a thermoforming protocol was established using a slow temperature ramp of 4°C per minute from room temperature to 160°C, a dwelling time of 60 min at 160°C, and a cooling step with a slow ramp to cool down until room temperature was reached. Then, the 3D flexible *kirigami* MEA and the lower mold were separated using droplets of water to facilitate the separation.

### 2.2. Variants, limitations, and yield of the kirigami-based process

Our *kirigami*-based process allows the facile customization of 3D flexible MEAs with different laminar and spatial requirements, thus enabling the implementation of designs that meet the spatial resolution for a large range of different neural applications. For example, 3D *kirigami* MEAs with shank lengths up to 225 µm (*Kiri*-225, Figure 2A) are ideal for recording from within the laminar structure of neural slices, such as explanted retinas or brain slices, while larger shank lengths up to 500 µm (*Kiri*-500, Figure 2B) or 1 mm (*Kiri*-1000, Figure 2C) provide access to deeper cortical layers in the intact rodent brain. While shanks longer than 1 mm were not tested, self-buckling calculations revealed that when maintaining the same cross-section of the PaC shank, a critical length of *Lc* = 20.6 mm is theoretically feasible. Hence, to enable intraneural access, each shank contained multiple electrode sites, and to interact with the surface of nervous tissues, we incorporated surface electrodes on the planar surface of the 2D *kirigami* template.

**Figure 2:**
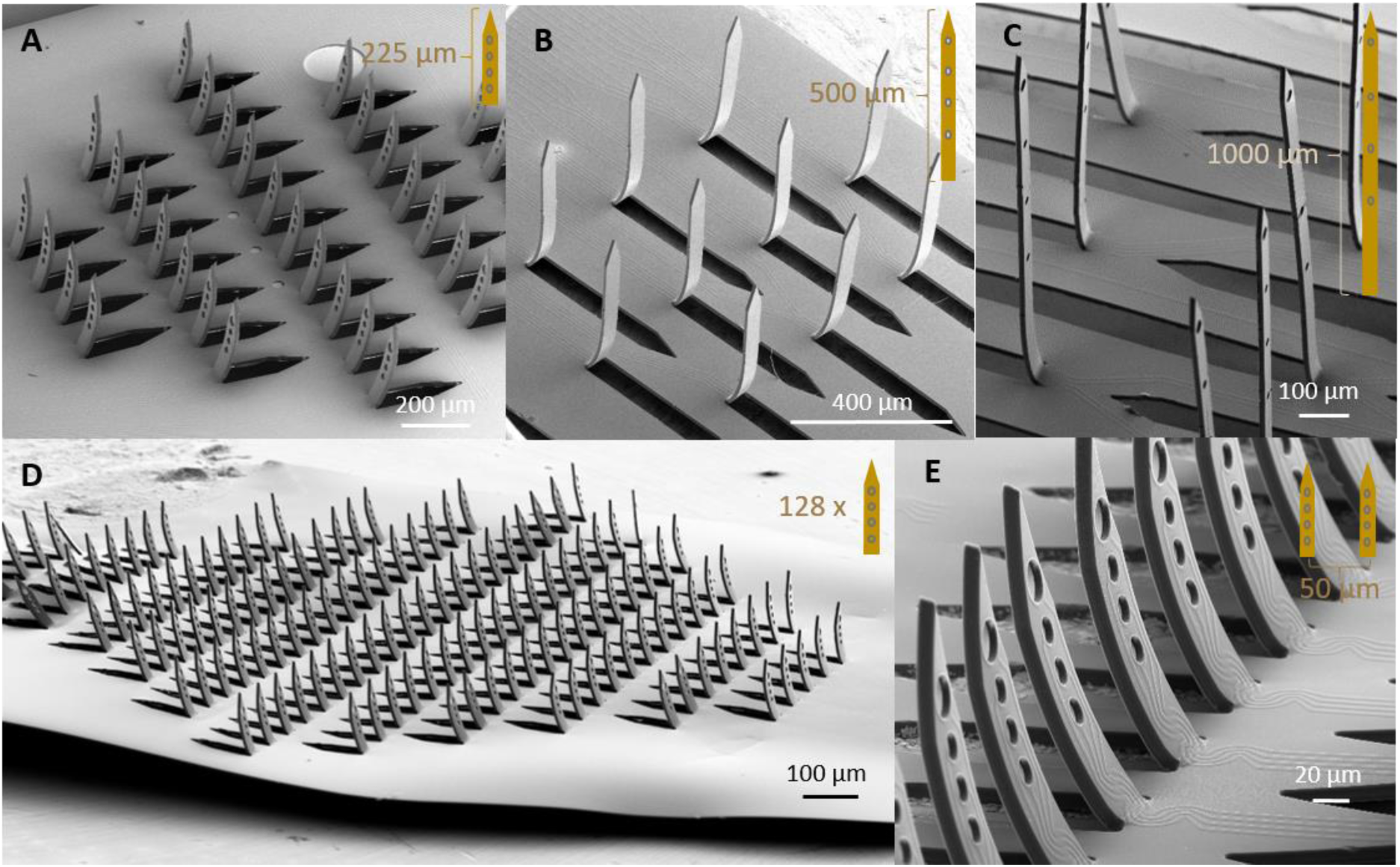
Variants of the 3D *kirigami* MEA design. Depending on the intended application, individual shank heights are feasible: from 225 µm for the retina and brain slices (A), over 500 µm for the cortex of rodents (B), to so far up to 1000 µm (C). Exemplary 3D *kirigami* MEA with up to 128 shanks on one flexible probe (D) and an inter-shank distance as low as 50 µm (E).

Additionally, the protruding structures of both the lower and upper molds must be mechanically stable to avoid self-buckling if straight shanks are desired. This stability depends on the geometry and dimensions of the protruding structures, as well as the mechanical properties of the mold materials and the lithographic techniques employed. In this work, we used two-photon lithography to enable the 3D printing of structures with aspect ratios (shank length/width) of up to 16.3. Calculations for self-buckling in the mold designs indicated that the *L_c_* for the protruding blocks is at least 61.9 mm. This exceeds the maximum printing range of the tool used (max. height: 8 mm), suggesting a potential limit to producing longer shanks. Furthermore, given that the molds can be fabricated with any additive manufacturing process that allows µm-mm resolution (*e.g.*, digital light processing printers or projection micro stereolithography^37^), and that 3D designs are derived from 2D templates, customizability in the design of the array of shanks is an advantage. In this work, we tested shank arrays with parallel rows and diamond-shaped designs (Figure S3).

Exploring the shank folding scalability of the matched die forming process, devices with up to 128 shanks, each 50 µm wide and 225 µm long and an inter-shank spacing of 100 µm were successfully fabricated (Figure 2D). To increase the electrode count, each shank contained four electrodes for a total of 512 electrodes. Two strategies would allow further increasing the electrode count without increasing processing time. The first strategy involves reducing the width of the feedlines and the diameter of electrodes, implying a geometrical constraint given by the resolution limits of the lithographic methods used. By optimizing the lithography process, it is possible to calculate the number of feedlines and electrodes that fit not only within a single shank but also the available area in the 2D *kirigami* template, while also maintaining a three-layered sandwich with a single metal stack layer. In our case, feedlines down to 3 µm and electrodes up to 25 µm were implemented with maskless lithography. Nonetheless, submicron features are also achievable with other techniques, such as electron-beam lithography as reported by others^38^. A second strategy consists of increasing the shank density; however, the success of the mechanical pressing is limited by the remaining solid area in the 2D *kirigami* template, especially the inter-shank spacing. The solid areas serve as mechanical support during the matched die compression. Therefore, the closer the shanks, the smaller the inter-shank area becomes, and the higher the compressive stresses, leading to structural instability by ripping the 2D template (Figure S4). To further explore this limitation, a *Kiri*-225 probe with an inter-shank spacing of 50 µm was successfully folded (Figure 2E).

Considering the design requirements described above, our *kirigami* folding process showed a successful folding yield of 98% (N = 100 devices including all designs). Successful folding was defined as the probe being separated from the molds without any difficulties (*e.g.,* without sticking to the mold), and all shanks being folded upright. After thermoforming, the 3D *kirigami* MEAs were coated with PEDOT:PSS *via* chronoamperometry to enhance the electrochemical properties of the base Au electrodes. After electrodeposition, we achieved a yield of 47% ± 16% for successful PEDOT:PSS deposition in different runs of the 2D fabrication process (N = 1721 electrodes distributed on 100 probes, and 7 fabrication runs). The high standard deviation and optical inspection indicate that the success of the process depends heavily on the accurate handling of the 2D *kirigami* template as well as on the quality of the underlying metal layer. Although the folding process has the potential to be carried out in an automated manner with *e.g.*, pick and place techniques and the use of micromanipulators, the alignment of the 2D *kirigami* template with the corresponding molds and the folding of the probes in this work were carried out manually. Hence, a learning curve was observed as the yield was lower in the first fabrication compared to the last run. When handling the 2D *kirigami* template during folding, hectic movements during the alignment can lead to cracks in the PaC layer, which can extend to the metal layer. Considering this learning curve, we achieved a yield of 80 % (N = 271 electrodes, last fabrication run) after the electrodeposition process.

### 2.3. Performance of *3D* kirigami MEAs

#### Mechanical considerations

When folding the 2D *kirigami* template, the PaC-metal-PaC sandwich that composes the implant is exposed to mechanical stress. When a shank bends upwards, the top surface compresses while the lower surface stretches (Figure 3A). Accordingly, finite element method simulations in COMSOL showed a maximum of -2.3 % volumetric strain at the center of the kink region on the top side and 1.2 % on the bottom side. After mechanical testing, we determined the Young’s modulus of the 10 µm-thick PaC layer thermoformed at 160 °C to be 1.70 GPa ± 0.32 GPa. Consequently, we reached a strain of up to 13.93 % ± 11.25 % at break. Despite the high standard deviation in the measurements, a maximum strain of 2.3 % during folding does not mechanically alter the PaC layer as it is still within the linear (elastic) region of the material. Hence, assuming a homogenous volumetric strain distribution, the metal layer at the middle of the two 5 µm-thick PaC layers is exposed to 0.55 % strain, which is not damaging the Au^39^. After folding, SEM inspection confirmed a PaC-Metal-PaC sandwich structure with PaC wrinkling at the kink area of the shanks (Figure 2). This observation supports the assumption that the feedlines (straight as well as meander-shaped ones) were not damaged during the folding process.

**Figure 3:**
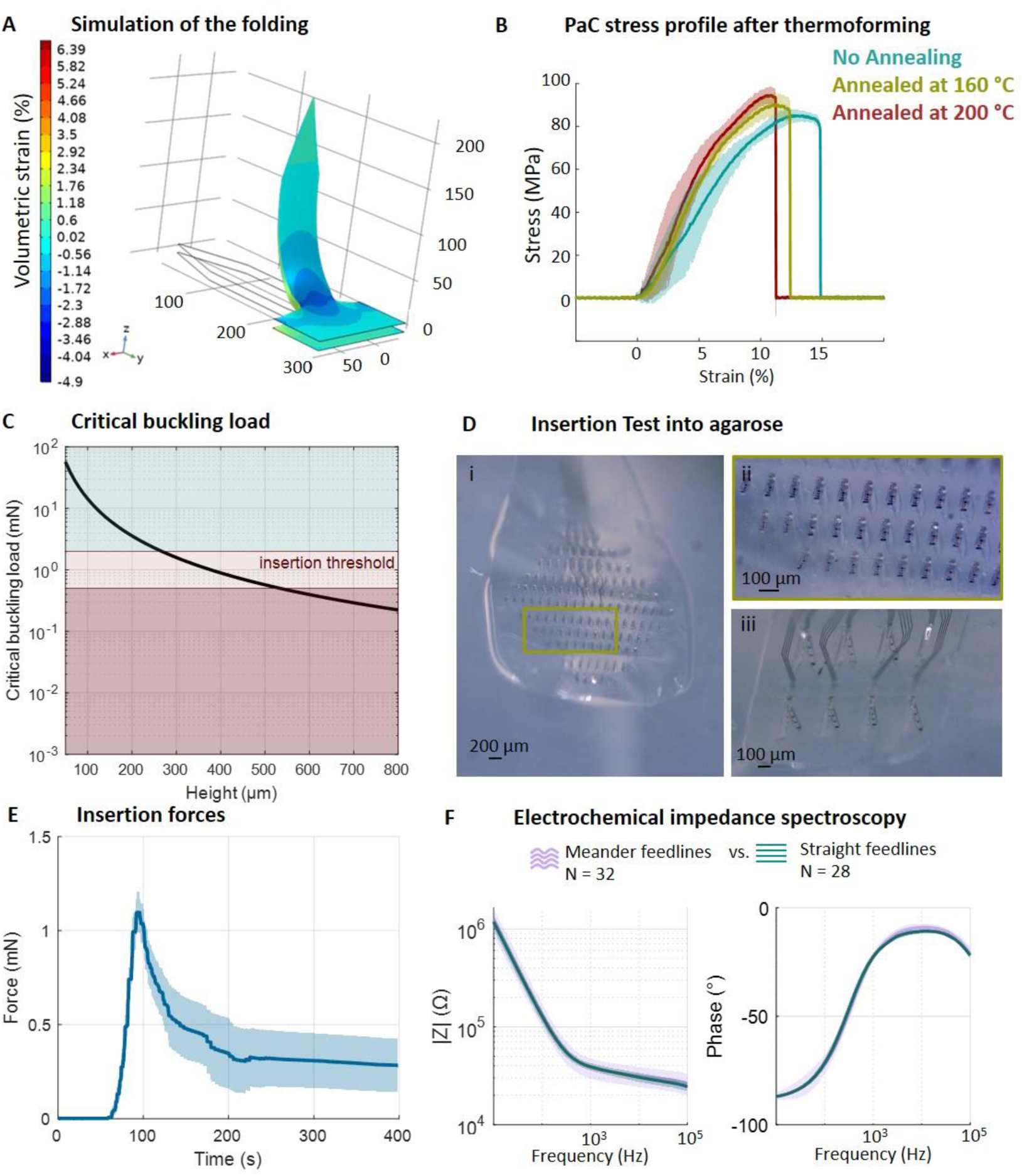
Mechanical and electrochemical performance of 3D *kirigami* MEAs. A) FEM analysis of the volumetric strain upon folding a flexible PaC shank at a 90° angle. B) Stress – strain (time) curve of 10 µm PaC films untreated (blue), annealed at 160°C (green) and annealed at 200°C (red). C) Theoretical buckling load of a 10 µm thick and 50 µm wide PaC slender column of different heights. The insertion threshold according to the literature (0.5 to 2 mN) is marked in red. D) Insertion of a *Kiri*-225 with 128 shanks (D_i_) and zoom-in (D_ii_), and insertion of a *Kiri*-500 with eight shanks into tissue phantom (agarose gel) (D_iii_). E) Insertion force profile of a *Kiri*-500 with 8 shanks (N = 7) penetrating agarose tissue phantom. F) Bode plot of the electrochemical impedance spectra of electrodes with a diameter of 15 µm located in PaC shanks with meander or straight feedlines at the kink (mean ± standard deviation).

To determine an optimal thermoforming protocol, we analyzed the mechanical properties of flexible PaC exposed to thermoforming processes using maximum temperatures of 160 °C or 200 °C and compared the results to untreated samples (Figure 3B and Table S1). For 200 °C the Young’s modulus as well as stress and strain at break are statistically different in comparison to untreated PaC stripes. In turn, thermoforming at 160°C does not significantly change the Young’s modulus. Thus, we selected 160°C as the optimal thermoforming temperature.

Additionally, we evaluated the mechanical stability of the flexible 3D *kirigami* MEAs by predicting the critical buckling load of individual shanks to assess their penetration capability into neural tissue. To penetrate neural tissue, an insertion force *F_i_* between 0.5 and 2 mN is needed^40^. If *F_i_* exceeds the critical buckling load of the shanks, they fail due to bending upon tissue penetration. Thus, we predicted the insertion likelihood by calculating the Euler’s critical buckling load of a 10 µm-thick PaC layer stack thermoformed at 160 °C (Figure 3C). This showed that a maximal insertion force of 0.5 mN enables the insertion of shanks with a length up to 530 µm into neural tissues, *e.g.*, the brain without meningeal layers such as the dura, without using an additional insertion aid.

Prior to functional testing in neural tissues, we evaluated the insertion performance of our *kirigami* probes with agarose tissue phantoms, demonstrating the successful insertion of a *Kiri*-225 with 128 shanks and a *Kiri*-500 with eight shanks (Figure 3D, Supplementary Video SV2). In the latter case, the insertion of eight shanks resulted in a peak force of 1.1 ± 0.2 mN (N = 7 devices) (Figure 3E), which exceeds Euler’s critical buckling load of 0.57 mN for a *Kiri*-500 shank. However, since the measured peak force reflects the combined load across all the shanks, the *Fi* per individual shank is expected to be lower, supporting the successful insertion predicted in Figure 3C. As a result, *Kiri-*500 probes were selected for later testing *in vivo* where longer shank lengths were required.

#### Electrochemical characterization

A PEDOT:PSS coating was electrochemically deposited on the Au electrodes to improve the electrochemical performance of the 3D *kirigami* MEAs. An average impedance of 39.1 ± 3.4 kΩ at 1 kHz and 1.3 MΩ ± 0.2 MΩ at 10 Hz was measured for straight feedlines for electrodes 15 µm in diameter. For meander-shaped feedlines, an average impedance of 38.4 ± 9.0 kΩ at 1 kHz and 1.4 ± 4.1 MΩ at 10 Hz were determined respectively. While the mean impedance values show only a slight difference, the higher standard deviation in meander-shaped feedlines suggests that the fabrication process for straight feedlines is more consistent. Therefore, straight feedlines were preferred for further experiments. The greater impedance variation in meander-shaped feedlines is likely due to their closer proximity to the edges of the shanks, which increases their susceptibility to minor cracks, which are a direct result of the RIE processing stage, as previously discussed.

Accordingly, the electrochemical performance of the electrodes shows the expected resistive behavior of PEDOT:PSS electrodes at the high frequency band (above 1 kHz), relevant for recording action potentials. At low frequencies (below 100 Hz), where LFPs are usually measured, the electrodes exhibit a capacitive behavior with the phase angle approaching -90° ^41–43^. Hence, 3D *kirigami* MEAs showed optimal characteristics for electrophysiological recordings, as low impedance values improve the recording quality of neuroelectronic interfaces^44^.

### 2.4. Applications of 3D kirigami probes

To highlight the utility of the 3D *kirigami* probes for characterizing both pathological and healthy activity across various regions in the central nervous system, and showcase their versatility in capturing neural dynamics at different spatial scales, we deployed customized versions for different applications *in vitro* and in *vivo*. We used *Kiri*-225 probes in human brain slices to study *in vitro* epileptic models, and *Kiri*-500 probes to map somatosensory and visual stimuli in acute and chronic settings in the intact brains of anesthetized and awake mice, respectively.

#### Explanted human brain slices

*Kiri*-225 probes were first utilized to examine the spatiotemporal propagation of epileptic activity in acute human brain slices and organotypic cultures. The brain slices were obtained from recessed access cortical tissue of epileptic patients at the University Hospital Aachen. As previously reported, seizure-like events (SLE) can be triggered by pharmacological manipulation^45^. An SLE is defined by three key characteristics: a paroxysmal change in neural activity that is distinct from the background activity, a temporal and spectral evolution of discharges, and a return to baseline activity^46,47^. During SLEs, ictal discharges (IDs) are observed, which contain Delta and Gamma waves with increased spectral power in a frequency range between 1-100 Hz , high-frequency oscillations (HFOs) in the 250-350 Hz range, and increased multi-unit activity (MUA) above 300 Hz^45^. As the thickness of explanted human brain slices ranged from 250 to 300 µm, *Kiri*-225 probes incorporating both surface and penetrating electrodes were utilized for electrophysiological recordings.

Human brain slices were submerged in artificial cerebrospinal fluid (aCSF) and placed in a perfusion chamber (Figure 4A). To insert the *Kiri*-225 probe, we first positioned the device on the brain slice surface using a micromanipulator and then gently pressed it into the tissue with a metal rod attached to a second micromanipulator (Supplementary Figure S5A). The probe was inserted such that individual shanks spanned multiple cortical layers. This placement was confirmed in two samples through immunohistochemical staining, which demonstrated the position of the shanks across cortical layers L2/3 to L5/6 (Figure 4B, Supplementary Figure S5B). This placement allowed for recordings of neural activity at different tissue depths and cortical layers. Due to the uneven surface of the brain slice and the slightly smaller surface area of the metal rod compared to the probe (see Supplementary Figure S5A), some shanks may not have fully penetrated the tissue. Despite this, the electrical activity captured by the electrodes connected to the data acquisition system confirmed the successful insertion of the shanks located in the middle of the probe.

**Figure 4:**
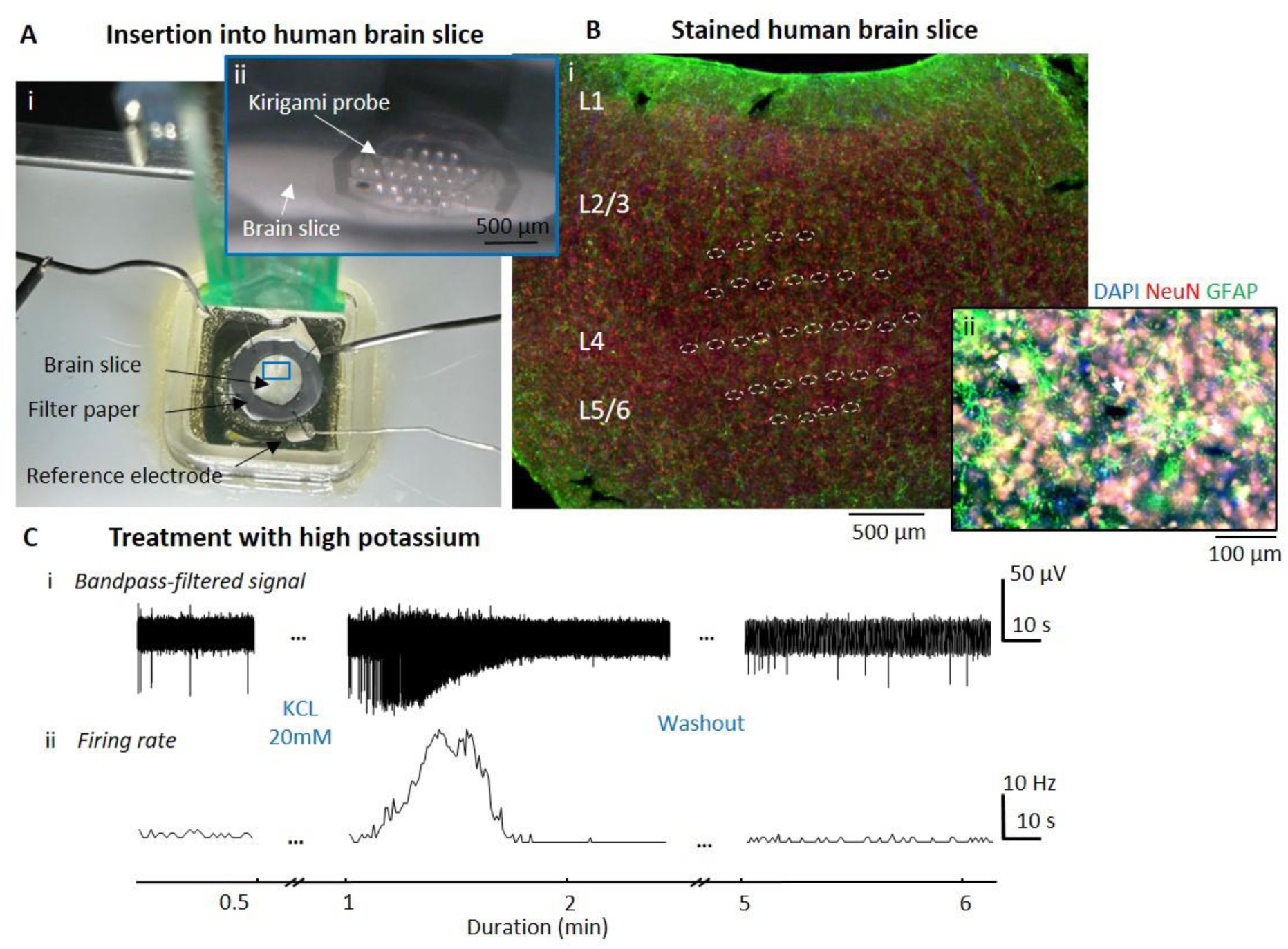
Electrophysiology with *in vitro* human brain slices. A) Insertion of a *kirigami* probe into a human brain slice inside a perfusion chamber with constant inflow and outflow of aCSF. The brain slice is submerged in aCSF and is supported by a doughnut-shaped filter paper and fixed on a PDMS supporting pillow with insect pins, while the *Kiri*-S probe is inserted perpendicular to the tissue. B) Immunohistological staining for NeuN (red), GFAP (green) and DAPI (blue) showing the insertion sites of a *Kiri*-225 probe. C) Human brain slice response to treatment with a high extracellular potassium concentration of 20 mM. Bandpass-filtered data (C_i_) and the firing rate along the complete recording (bin size = 1 ms) (C_ii_), show that the cells reach a depolarization block under the treatment and go back to baseline activity after washout.

We first confirmed that the electrical recordings indeed captured neural activity. To do so, we initially recorded spontaneous activity and then induced a physiological response by increasing the K^+^ concentration from 3 to 20 mM, which is known to cause a depolarization block. The latter was observed shortly after adjusting the K^+^ concentration to 20 mM, as the spiking activity strongly increased with larger spike amplitudes and higher firing rates (Figure 4C_i_ and 4C_ii_). Following the increase in activity, a depolarization block occurred, causing the complete cessation of subsequent action potentials. When K^+^ concentration was restored to normal levels, spontaneous activity quickly returned to its baseline state. This observation is consistent with the mechanisms of action potential formation and aligns with earlier findings^48,49^.

In a second set of experiments, we aimed to study the spread of epileptic activity within six human brain slices by adjusting the extracellular ion concentration to induce epileptic activity patterns. Following the protocol by Pallud *et al*.^45^, we modified the aCSF medium by increasing the K^+^ concentration to 8 mM and reducing the Mg^2+^ concentration to 0.25 mM. After introducing the modified aCSF medium, SLEs were observed. The duration of SLEs was determined by a sustained increase in the firing rate of spiking activity (100 – 3000 Hz) that exceeded the baseline firing rate in each individual electrode. We observed SLEs in five out of six slices from three different patients with a signal-to-noise ratio (SNR) of up to 200.9 and mean of 6.0 +/-1.6. On average, an increased firing rate during SLEs lasted 51.4 s, with a standard deviation of 73.5 s (*N* = 88 SLEs). These events occurred repeatedly, with intervals averaging 144.2 ± 170 s. Initial inter-SLE periods were as long as 809 s but decreased over time to as short as 6 s (Figure 5A). SLEs were observed after an average of 33.7 ± 13.7 min of perfusing the modified aCSF across six slices. During the SLEs, frequencies below 1 kHz predominantly contributed to the recorded electrophysiological signals, as evidenced by the increased spectral power (Figure 5A_ii_). This observation is consistent with previously reported studies^45–47,50^. Alongside the increased firing rates, HFOs (Figure 5A_iii_) were observed during the SLEs. Typically, the firing rate was highest at the beginning of the SLE and subsequently flattened until the end (Figure 5A_iv_). The average firing rate during a SLE was 7.8 ± 5.2 Hz, which was significantly higher than the overall average firing rate of 0.7 ± 0.7 Hz (Figure 5C). Furthermore, characteristic discharges during SLEs including HFOs were observed, exhibiting features of epileptic activity that matched temporal and spectral characteristics described in earlier studies^45^ (Figure 5B).

**Figure 5:**
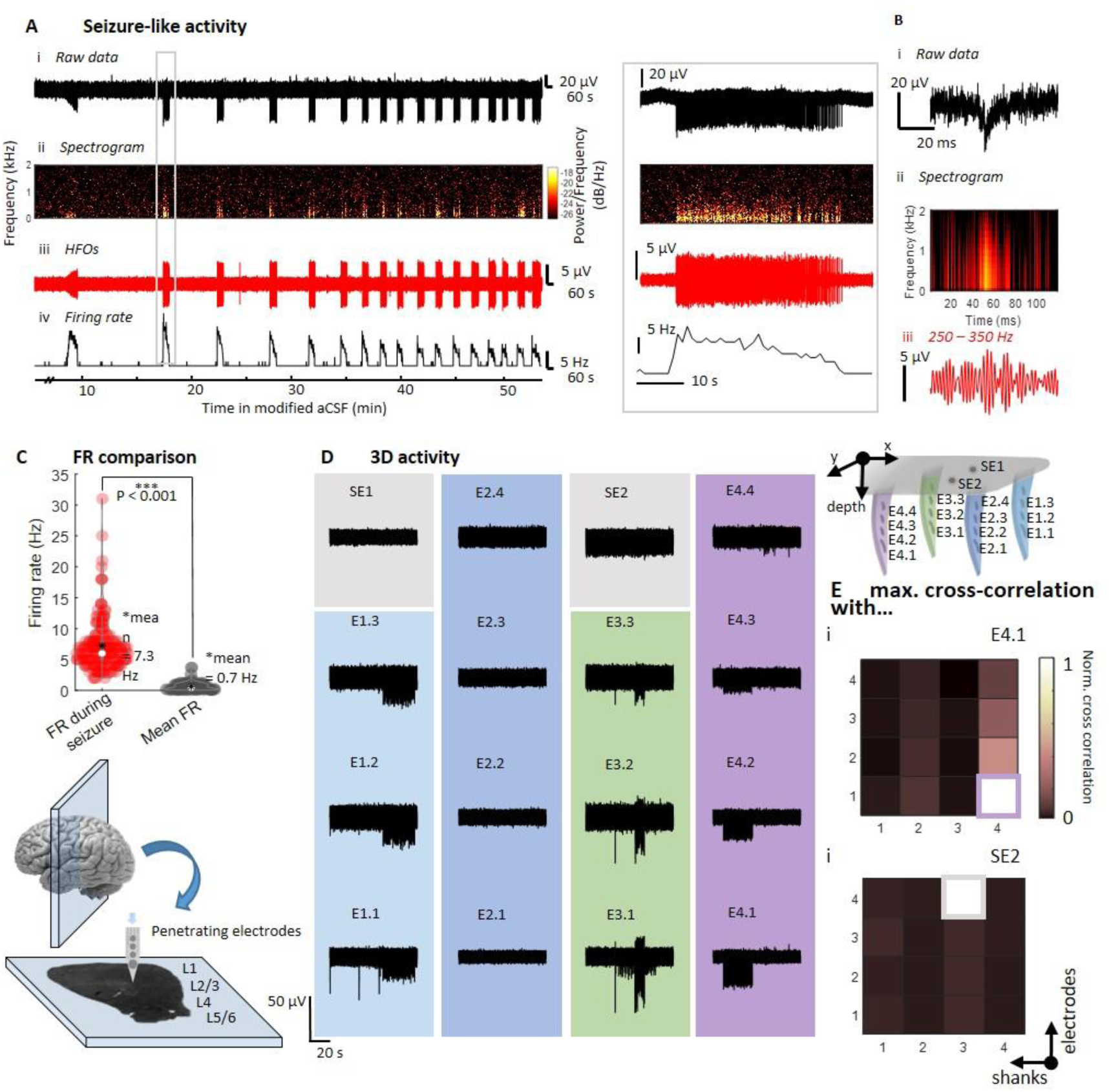
Seizure-like activity in human brain slices. A) Electrophysiological activity upon treatment with modified aCSF (8 mM K^+^ and 0.25 mM Mg^2+^), raw data (Ai), spectrogram (Aii), HFOs filtered at 250 – 350 Hz (Aiii) and firing rate traces (Aiv). B) Examples of an ictal-like discharge, raw data (Bi), spectrogram (Bii), and filtered HFOs (Biii). C) Comparison of firing rates during SLEs and averaged over the whole trace (p < 0.001 using a paired t-test with N = 153 SLEs). E) 3D map of firing rate upon treatment with modified aCSF. The electrodes of four shanks (dark blue, light blue, green, violet) and the surface electrodes (grey) each capture SLEs at different time points. F) Maximal value of the cross-correlation for electrode E4.1 and SE2 with all the other electrodes.

Leveraging the spatial distribution of electrodes in the *Kiri*-225 probe, we then compared recordings from surface electrodes with those from penetrating electrodes. This analysis revealed the presence of ictal-like discharges that were detected by electrodes located inside the neural slice, allowing for the identification of the onset of SLEs across electrodes on individual shanks. Figure 5D depicts a representative recording where the onset of an SLE (shank 4, pink column) occurred at approximately 160 µm (centered on a 25 µm diameter recording electrode) below the tissue surface. The SLE spreads vertically over about 60 µm, based on the center-to-center distance, between E4.1 and E4.3 but is not visible in the surface electrode. This further demonstrates the capability of penetrating 3D *kirigami* MEAs to detect SLEs within the tissue parenchyma that would otherwise be missed by surface or distant electrodes (*e.g.*, SE2 or E4.4).

Furthermore, as illustrated in Figure 5D-E, SLEs captured across individual shanks separated by 100 µm and 300 µm in *x* and *y* axis, respectively, were largely independent from each other. Cross-correlation analysis of the data recorded with individual electrodes and all other electrodes demonstrates that electrodes within a single shank, such as shank 4 (Figure 5Ei), exhibited a high degree of correlation, whereas the cross-correlation with neighboring shanks was low. Similarly, the surface electrode SE2 was not significantly correlated with any of the penetrating electrodes, nor with the other surface electrode SE1 (Figure 5Eii). These findings indicate that the epileptic events captured within the local neural networks remained in the vicinity of the individual shanks instead of spreading across all cortical layers. This behavior was observed in all the recorded slices.

Our *in vitro* recordings with 3D *kirigami* MEAs provide evidence of the occurrence of isolated SLEs at different time points and depths within human brain slices. These findings underscore the importance of studying neural activity patterns in brain slices not only with 2D MEAs on the tissue surface but through 3D volumetric sampling. The slicing procedure can particularly damage cells at the tissue surface. Therefore, in contrast to planar MEA systems, the penetrating *Kiri*-225 probes offer a significant advantage by incorporating both surface and penetrating electrodes. This configuration allows for the recording of neural activity not only on the surface of the tissue but also within the slice parenchyma and throughout the tissue volume. The utilization of 3D *kirigami* MEAs in human brain slices offers a more comprehensive understanding of pathological activity patterns by localizing the spatial origin of SLEs, tracking their spread through localized networks, and simultaneously observing neural activity from other local networks.

#### In vivo recordings from mouse cortex

Aside from applications *in vitro*, flexible 3D *kirigami* MEAs are promising tools for brain recordings *in vivo*, especially in the cortex. Because the design and electrode density of *kirigami* probes can be flexibly adjusted to match the spatial layout of different brain regions, this allows custom applications that require observing the spread of neural activity across different cortical layers and brain areas. Such recordings across layers and regions could be highly valuable for various clinical applications, for example to study the volumetric spread of epileptic activity throughout the cortex or for chronic BMIs to capture rich sensorimotor signals from local neural networks.

To test the *kirigami* probes for *in vivo* applications, we first implanted them in the somatosensory cortex of anesthetized mice. The somatosensory cortex is composed of specialized regions for different body parts^51^, which makes it possible to relate their respective neural activity to tactile stimulation of the hindlimb and identify region- and layer-specific functional responses (Figure 6A-D). To target both superficial and deeper cortical layers in different regions, we implanted *Kiri*-500 probes with eight or ten shanks. Before the implantation (Figure 6A), we used cortical widefield imaging to identify the hindlimb region of the primary somatosensory cortex (Figure 6B). Mice were transgenic animals that expressed the calcium indicator GCaMP6s in all excitatory neurons^52^, allowing us to measure changes in neural activity as changes in fluorescence light. The somatosensory hindlimb area was then identified as the region with the strongest average neural response to brief deflections of the left hindlimb under isoflurane anesthesia (*n* = 50 trials). To obtain a 3D recording from multiple cortical regions *in vivo*, we then placed the shanks of the probe within the somatosensory hindlimb region, using the border of the widefield response as a guide. For this purpose, a high-resolution vessel image (Figure 6B_i_) was used to identify the accurate location for probe placement.

**Figure 6:**
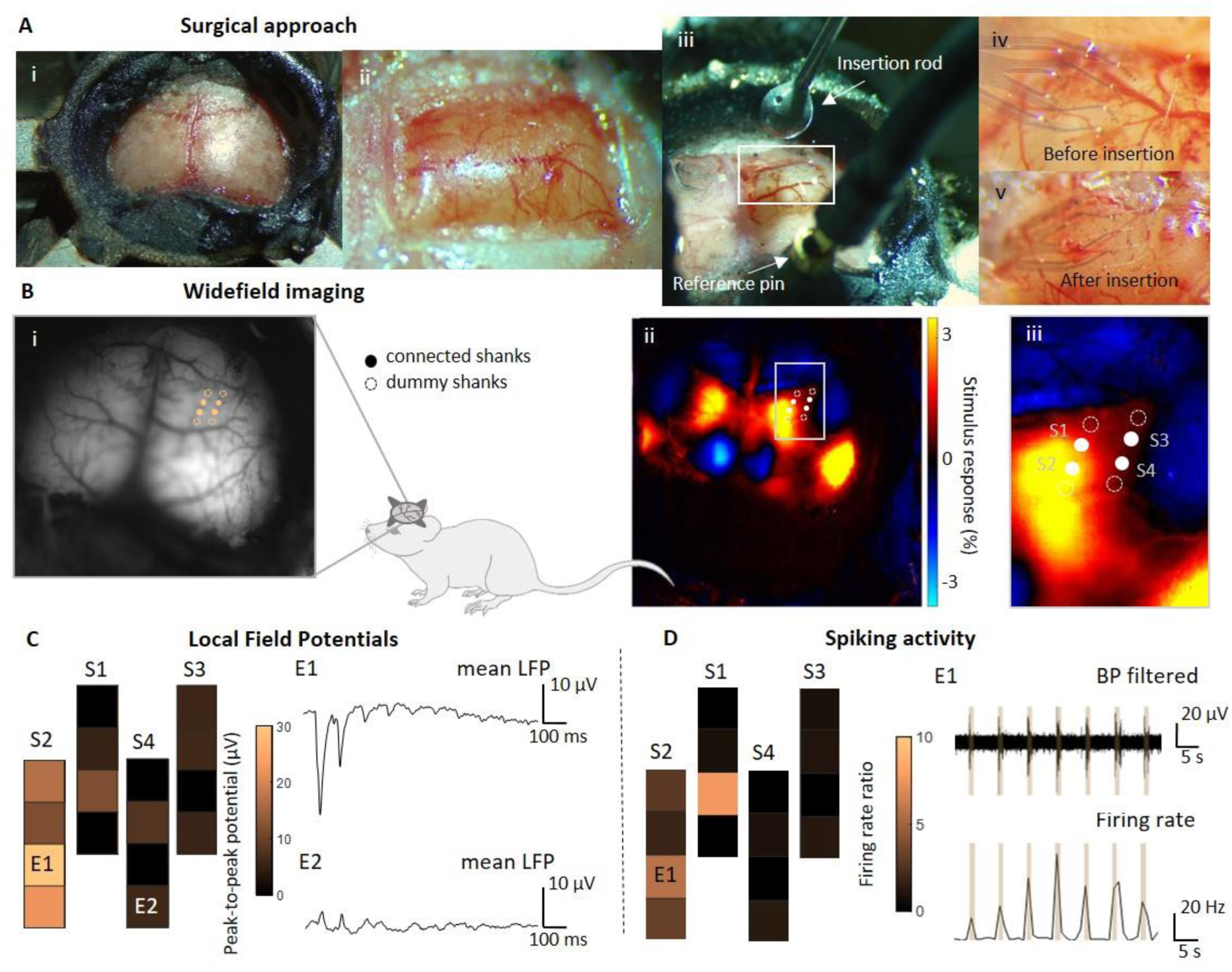
3D electrophysiological recordings of acute *in vivo* mouse somatosensory cortex. A) Surgical approach: A head bar is fixed on the mouse head (i) before the craniotomy (ii). The implant is pushed inside the cortex using an insertion rod (iii). The shanks are therefore placed at the desired position (iv). After the retraction of the insertion rod, they remain at their position (v). B) To identify the optimal positioning of the shanks across different brain regions (i), widefield imaging was used to identify cortical responses to tactile hindlimb stimulation (ii and zoom-in iii). C) Averaged LFPs show the spatial activity spread depending on the depth (z) and x-y location of the electrodes. The peak-to-peak potentials were based on the averaged LFPs over 50 trials. D) Average changes in the spiking activity equally differs according to the electrodes position. Shown are changes in firing rates due to tactile stimulation. The firing rates rise respectively with the stimulation (brown bars).

Following the addition of a head bar (Figure 6A_i_), a craniotomy was placed (Figure 6A_ii_) and the dura was removed, allowing the probe shanks to be inserted at the desired position of the somatosensory cortex. Initially, a micromanipulator was employed to position the implant on the cortical surface. To facilitate the insertion of the implant into the tissue, a rod was attached to a second micromanipulator and placed directly over the implant (Figure 6A_iii_). After careful positioning, all shanks were simultaneously inserted by slowly moving the micromanipulator downwards. This allowed the shanks to overcome tissue dimpling and seamlessly penetrate the cortex without any visible tissue disturbance (Supplementary Video SV3). Prior to insertion (Figure 6A_iv_), the shanks were visible on the surface of the cortex. Following insertion (Figure 6_v_), the insertion rod could be retracted while the shanks remained stably inserted in the tissue. After the insertion, we repeatedly produced tactile stimuli with a custom-made stimulator, producing 100-ms-long deflections of the hindlimb with a small servomotor.

Neural responses to tactile stimulation were clearly visible as LFP deflections (signals below 100 Hz) (Figure 6C, S6) and differed across the cortical locations of each shank. For each electrode, we computed the average LFP response to tactile stimulation and their peak-to-peak potentials. In agreement with our widefield imaging results, the neural response to tactile stimulation in the two shanks (S1 and S2) within the hindlimb region of the somatosensory cortex was much stronger compared to the lateral shanks S3 and S4 that were placed outside the hindlimb region. The strongest response occurred in electrode E1, located closest to the hindlimb region center at a depth of 300 µm, exhibiting an average potential of 28 µV, whereas electrode E2 on the neighboring shank S3 only yielded an average potential of 5.6 µV.

The analysis of the high-pass filtered signals (100 to 3000 Hz) to isolate neural spiking activity revealed similar results (Figure 6D, S6). During the stimulation period, the firing rates mostly increased in electrodes near neuronal units that responded most strongly to tactile stimulation. Again, the electrodes on the S1 shank, such as electrode E1, showed a strong, stimulus-induced change in firing rates. This was also the case for electrodes on shank S2 whereas few changes in firing rates occurred on shanks S3 and S4. In general, spiking activity was easier to capture at deeper cortical layers, as cortical layer 1 at the surface of the cortex is mainly composed of axons. Consequently, while high amplitude LFPs were captured with all four electrodes of S1, high spiking activity was mostly captured with deeper implanted electrodes, such as E1.

Our results demonstrate the capability of the *Kiri*-500 probes to record neural responses across layers and cortical regions in the living brain. To extend this further, we also used *Kiri*-500 probes in a chronic preparation with awake mice. Here, we recorded neural responses to different visual stimuli across different depths and cortical areas to characterize sensory neural responses in the awake brain. After the insertion, we placed a glass window on top of the implant (Figure 7A_i_) and secured it with dental cement to the skull (Figure 7A_ii_). This allowed us to stabilize the implant in the cortex and monitor its position through the glass window (Figure 7A_iii_). For this chronic application, the front-end connector of the implant was customized for a total form factor of 21 x 11.5 x 7.5 mm and weight of 1.6 g, being small and light enough for the mouse to carry (Figure 7A_iv_). Awake mice were then placed inside an experimental setup (Figure 7Av) with a running wheel and a monitor displaying visual stimuli that fully covered the visual field of the right eye, contralateral to the implanted cortical hemisphere. To functionally determine the location of visual cortical areas before the implantation, we used widefield imaging and visual retinotopic mapping and identified the location of the primary cortex and several higher visual areas (Figure 7b_i_, blue and red regions). The functional results were in good agreement with the expected location of visual areas (white outlines), therefore we placed the *Kiri*-500 probe to cover the primary visual cortex (V1) and several higher visual areas.

**Figure 7:**
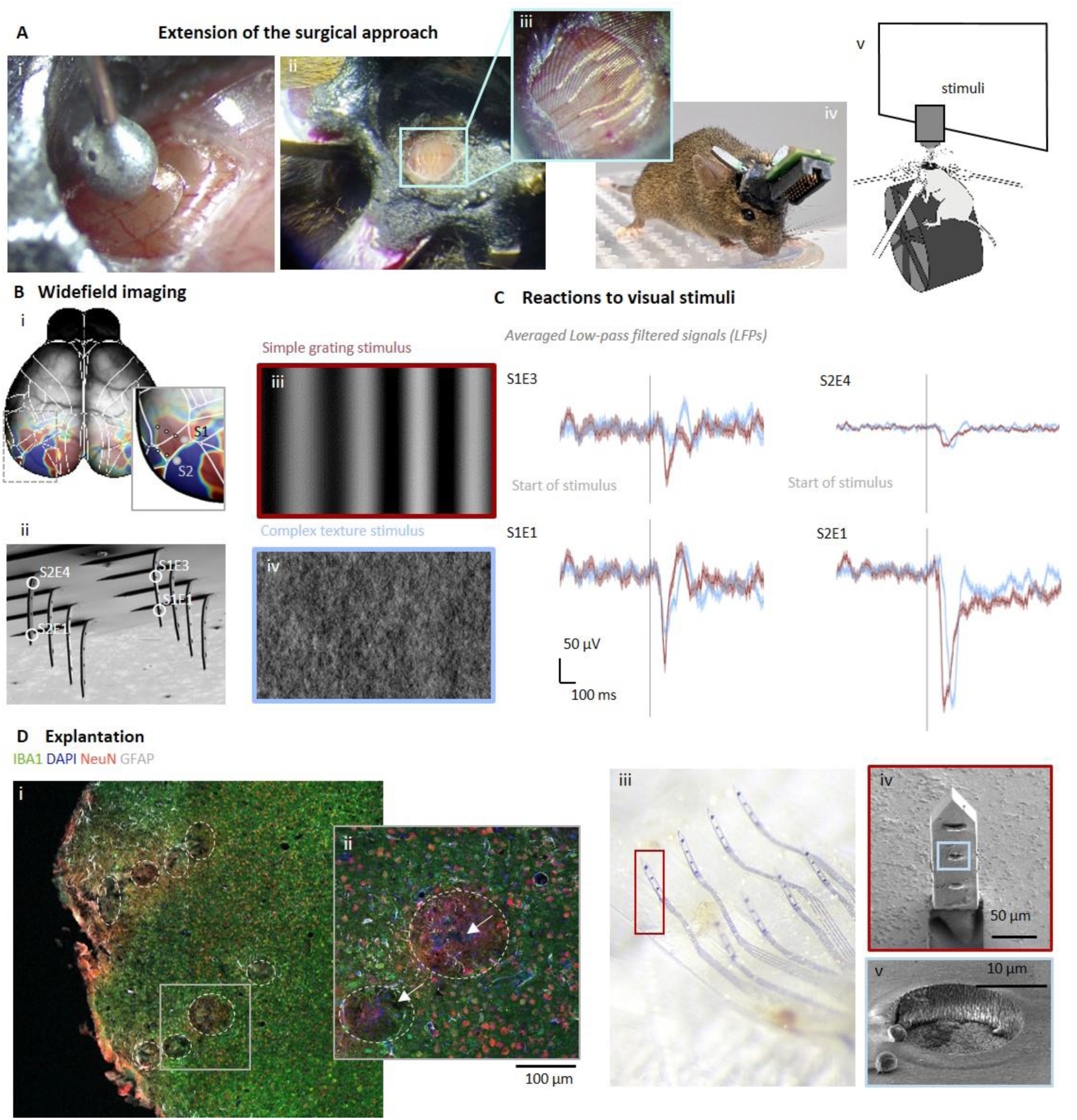
Chronic implantation and recordings from the visual cortex of mice. For visual stimulation with awake mice, the surgical approach from Figure 6 was extended (A): The insertion rod to place a glass window on top of the implant (Ai, Aii and zoom-in Aiii). After recovery time, the mouse was woken up (Aiv) and placed inside the experiment setup (Av) where it was exposed to visual stimuli. Widefield imaging was used to determine the desired positioning of the shanks (Bi-ii) upon visual stimulation with simple grating (Biii) and complex texture (Biv) stimuli. Visually evoked averaged LFP responses in shank 1 and 2 (C). Immunohistological stainings for mature microglia (IBA1, green), cell nuclei (DAPI, blue), astrocytes (GFAP, grey) and mature neurons (NeuN, red) of 20 µm-thick horizontal slices after brain perfusion and the retraction of the implant (D) showing implantation lesions with marked large ROI (white). Additionally, neurons expressing GCaMP6 are also green (Di). Dii shows one hole tracked through different depths and the marked large ROI (white). Implantation lesions (small ROI) match the dimensions of the shanks (white arrow pointing to ∼50 µm wide hole) and the implantation locations of the shanks. Optical inspection of the explant with light microscopy (Diii) and SEM (Div,v) reveals that the shanks of the *Kiri*-500 implant were still folded upwards and adsorption of biological matter on the electrodes (Dv).

We then characterized how different cortical areas and layers responded to two types of visual stimulation: a simple moving grating stimulus and a complex texture (Figure 7Biii and 7Biv). Focusing on LFP responses in two neighboring shanks (S1 and S2), we found clear differences in the average stimulus response across depths and locations (Figure 7B-C). Electrodes S1E1 and S2E1, positioned in the deeper cortical layers 2/3, exhibited stronger visual responses compared to the superficial electrodes S1E3 and S2E4 in layer 1. This difference in response was particularly pronounced in V1 (shank S2), whereas in the more medial anterior cortex (area AM, shank S1), the responses were more consistent across different depths. Moreover, while both gratings and texture stimuli induced strong neural responses in V1, visual responses in AM were weaker and more selective for gratings over texture stimuli, especially in the superficial electrode S1E3 (red versus blue traces). Interestingly, we also observed a time shift in the response delay to visual stimuli in V1, with textures eliciting a longer delay compared to gratings. This is likely due to the increased complexity of the texture stimulus, which requires more extensive neural processing in the visual system^53^. Together, these results show clear differences in the processing of complex visual stimuli across cortical depth and regions, demonstrating the suitability of *Kiri*-500 probes for chronic recordings in the awake brain.

Four weeks after implantation, the *Kiri*-500 probes were explanted, followed by immunohistochemical analysis of the implanted brain tissue (Figures 7D_i__-iv_, S7, S8) and optical inspection of the explanted electrodes (Figure 7D_ii_). 20-µm thick horizontal sections of the cortex were stained for mature neurons (NeuN, red), cell nuclei (DAPI, blue), astrocytes (GFAP, grey) and microglia (IBA1, green). The implantation footprint of the shanks is visible as small lesions (dark holes) referred to as small ROIs. Additionally, the slices reveal foreign body reactions (FBRs) in tissue regions in proximity to these small ROIs (Figure 7D_i_ and Figure 7D_ii,_ white circles). This FBR is evident as fluorescent changes in the surrounding tissue (large ROI), with small lesions (small ROIs) typically located midway and occasionally off-center within the affected areas (Figure 7D_ii_, white arrows). The insertion footprint, which contains multiple footprints of individual shanks, could be tracked throughout a depth of 260 µm. The mean area of the small ROIs, which was 478 ± 380 µm² (N = 24), correlates with the dimensions of the shanks. The mean size of the affected regions (large ROIs) exhibiting an FBR was 7467 ± 373 µm² (N = 56 large ROIs across 11 slices), which is approximately 15 times the cross-sectional area of the shanks (500 µm²).

Examination of the larger ROIs (FBR) across various depths were discernable until Z13, corresponding to an approximate depth of 260 µm, and exhibited a trend of larger footprints in superficial layers (layer 1) and smaller in deeper layers (layers 2/3) (Figure S8A). In contrast, the smaller ROIs were only visible until slice 10, corresponding to an insertion depth of approximately 200 µm (Figure S8B). Others reported similar observations^54^. Additionally, this can be explained by the fact that the shanks have a reduced cross-sectional area toward the tip, resulting in a smaller footprint in the deeper layers. Figure 7 D_ii_ shows the affected insertion area on slice 8, corresponding to a depth of 160 µm, with a reduction in astrocyte density near the insertion area (small ROIs). This was also demonstrated by analyzing the relative fluorescence intensity (Figure S8D), which also indicated a notable reduction in the number of astrocytes observed in the large ROIs (Figure S8C), similarly to previously documented studies^55^. In contrast, nuclei and neurons were not observed directly at the implantation site (Figure 7D_ii_ and 6D). Similar observations were documented in other studies^46^. The expression of the GCaMP indicator in excitatory neurons of the genetically modified mice, which also appear green, interfered with the ability to reliably analyse IBA1 staining.

However, it should be noted that the slicing was likely not perfectly perpendicular to the insertion direction, as achieving perfect alignment is very challenging. The total insertion depth may therefore have been underestimated, and the size of the ROIs overestimated when the slices were not cut perpendicular to the shank length. Additionally, pulling during implant retrieval caused irregularly shaped insertion holes, leading to considerable variability in the measured footprint areas (both large and small ROIs). Consequently, the assessment of FBRs is likely overestimated due to both the retraction of the implants and the slicing process.

Nonetheless, the shanks of the probe remained accurately folded upon implant retrieval (Figure 7D_iv_ and 7D_v_), and SEM inspection of the electrodes showed no evidence of delamination aside from the expected minor adsorption of biological matter. This highlights the long-term stability of the *kirigami* folding and electrodes of the shanks (Figure 7D_vii_). Together, the results of these experiments demonstrate the potential of our 3D *kirigami* probes to obtain recordings from different depths and brain areas inside living neural tissue, allowing for a detailed investigation of neural response patterns. Additionally, the results demonstrated the capability of the *kirigami* implants to withstand acute and chronic implantation *in vivo* while the amount of astrocyte accumulation as an indicator for FBR was low, especially in deeper layers of the cortex.

## 3. Conclusion and Outlook

This work presents a novel and scalable *kirigami* approach that enables the simultaneous folding of up to 128 shanks to create a flexible 3D MEA with both penetrating as well as surface electrodes. The dimensions and electrode counts of our *kirigami* MEAs surpass those of existing approaches for flexible 3D neural implants (Table S2) with a maximum shank length of 1000 µm, and a minimal inter-shank spacing of 50 µm. A variety of design configurations were investigated for potential use in diverse neural tissue applications and experimental settings, spanning from *in vitro* to *in vivo*. This demonstrates the customizability of our approach, which permits the configuration of diverse spatial requirements. Although the fabrication process remains manual, implants can be produced rapidly and reliably with a high yield. Our *kirigami* folding technique employs a matched die compression process, rendering it suitable for serial manufacturing. Nevertheless, the interconnection of all 512 electrodes remains a challenge due to the subsequent necessity of a considerable number of wires, which must be linked to a recording system. These components are typically costly and bulky. An ideal solution would be a wireless, high-channel connector design, which would be advantageous for long-term recordings. However, to the best of our knowledge, wireless front ends with 512 channels are currently not commercially available.

Our analysis of the mechanical characteristics showed that shanks with lengths of up to 500 µm, a width of 50 µm, and a thickness of 10 µm can penetrate neural tissue without buckling. However, to access deeper cortical layers, the cross-sectional dimensions of the shank must be adjusted to achieve a higher critical buckling load. This can be done by increasing the width or the thickness of the shank, enabling shuttle-free insertion into the tissue. Our 3D *kirigami* MEAs also feature a small cross-sectional footprint and flexible, biocompatible materials with low bending stiffness to promote optimal tissue integration. Through electrochemical impedance spectroscopy, we also confirmed low electrode impedances throughout the device, which are advantageous for high SNR recordings of spikes and LFPs.

Our study shows how flexible 3D *kirigami* MEAs enable capturing spatial dependencies in electrophysiological activity patterns in both *in vitro* and *in vivo* applications. The cortex, particularly in degenerated or diseased conditions such as epilepsy, is a complex system that requires further investigation to provide more effective diagnostic tools and treatments. Accordingly, a 3D MEA with high spatial and temporal resolution is highly beneficial for efficiently mapping electrophysiological activity throughout the 3D neural space. Nevertheless, the most advanced 3D neural implants are either merely recordings from specific x–y planes at designated z-depths within the neural tissue, as observed in Utah arrays^15,16^, or they are y–z neural threads that must be implanted in succession to create a comprehensive 3D mapping of the cortex^25–27^. In contrast, our flexible *kirigami* MEA, with a shank length of up to 500 µm and multiple electrodes for mapping multiple x-y-z locations, is implanted in a single step without the use of an insertion shuttle. This approach shortens the overall implantation and surgical periods and facilitates post-implantation monitoring of the implantation sites.

3D *kirigami* MEAs are therefore an ideal tool for advancing the understanding of neural information processing, particularly in relation to neurodegenerative diseases, such as seizure propagation in epileptic models. We successfully deployed the *kirigami* probes with penetrating and surface electrodes in healthy human brain slices *in vitro* and induced SLEs to examine their spatial origin and spread throughout the tissue. SLEs recorded in this study exhibited characteristics similar to those observed in previous studies, and the 3D sampling showed that SLEs occurred in separate local networks across cortical layers.

The complexity of neural networks in human brain slices *in vitro* and in the *in vivo* cortex of mice, especially in models of disease such as epilepsy, demand advanced 3D recording devices to study the volumetric propagation of electrophysiological activity^56,57^. SLEs in brain slices are often artificially evoked to allow a more detailed mechanistic manipulation that is not always feasible *in vivo*. While rhythmic neuronal activity is often associated with epileptiform events, it is important to note that various normal brain patterns also show synchronous bursts of activity. Surprisingly, if synchrony is measured between single cells and the network (measured at the population level), this is not always highest during the epileptiform event (*e.g.*, SLE), which reflects the complicated nature of epileptic activity^58^. In this study, we could also show that SLEs in human brain slices exhibit both synchronous and asynchronous components. These events spread within the 3D structure of the slices, highlighting aspects of SLEs that might not be detectable in traditional 2D MEA recordings^59^.

To demonstrate the utility of our 3D *kirigami* MEAs for volumetric neural recordings *in vivo*, we employed them in the somatosensory and visual cortex of mice. These sites were chosen because of their critical role in sensory restoration, such as touch or vision. We then aimed to demonstrate how spatial-dependent activity can be captured upon tactile and visual stimulation, exposing possible stimulation target sites for sensory restoration. In the somatosensory cortex, 3D electrophysiological activity was successfully recorded, as evidenced by the spatial-dependent LFP amplitudes and firing rates of individual neurons in response to hind limb stimulation. The differences in activity between neighboring shanks also accurately reflected their respective position in somatosensory subregions. Moreover, we found clear differences across cortical layers in individual shanks. In cortical layers 2/3, the incidence of spiking activity was higher in comparison to layers 1 and 4, underscoring the importance of 3D recordings to identify the most efficacious sites for electrical stimulation for the restoration of sensory functions.

Similarly, our recordings in the visual cortex of awake mice showed a clear spatial dependency upon different cortical layers. Furthermore, the responses exhibited variations across visual areas, depending on the stimulus complexity. After four weeks of implantation, a comprehensive examination was conducted on the electrodes and the implantation site. The folded *kirigami* shanks did not undergo any unfolding or delamination of the electrode material, demonstrating the long-term stability of the flexible 3D MEA structure. Immunohistochemical analysis of the tissue also showed that the cross-sectional footprint matched the dimensions of the *kirigami* shanks. Notably, an accumulation of astrocytes was observed only in layer 1, near the cortical surface and close to the insertion site. This suggests that the insertion impact was mild, likely due to the flexible and biocompatible properties of the *kirigami* implant.

From *in vitro* human brain slices to the *in vivo* cortex of mice, our experiments underscore the complexity of neural networks, particularly in diseased models where 3D recording platforms are a crucial tool to capture electrophysiological activity patterns across layers and brain regions. Our approach enables the straightforward fabrication of flexible 3D MEAs, capable of sampling the 3D architecture of neural tissues with highly customizable designs. This innovation combines the spatial sampling and resolution of devices such as the Utah and the Michigan array into a single, flexible neuroelectronic interface. With the ultimate goal in mind to better understand and consequently treat neurological disorders, such as epilepsy or loss of sensory functions, the flexible 3D *kirigami* MEAs presented in this work offer a highly valuable tool for future investigation.

## 4. Materials and Methods

### Fabrication of Kirigami 3D implants

#### Fabrication of 2D kirigami template

The fabrication of the 2D *kirigami* template comprised the interleaved deposition of two flexible thin film layers, one metal layer, and an electrode coating. All the steps are conducted in a certified cleanroom environment to ensure a stable fabrication^60^.

First, a 5 µm-thick PaC layer was deposited on a host silicon wafer *via* chemical vapor deposition using a PDS 2010 Labcoater 2 (Specialty Coating Systems Inc., USA). In a second step, a metallization process was performed using a lift-off technique. Here, the metal base layer for contact pads, feedlines, and electrodes was patterned after spin-coating the negative photoresist AZ LNR-003 (MicroChemicals GmbH, Germany) at 4000 rpm for 45 s with a ramp of 500 rpm/s, followed by a soft-bake at 120°C for 2 min on a direct contact hot plate. The photoresist was then exposed at 320 mJ/cm2 with a defoc of 2 and a CDB (critical dimension bias) of 800 with UV at 375 nm using a maskless aligner (MLA 150, Heidelberg Instruments, Germany), followed by a post-exposure bake at 100°C for 90 s on a direct contact hot plate, a developing step in AZ 326 MIF (MicroChemicals GmbH, Germany) for 90 s, and a cleaning step in deionized water. The wafer was then evaporated with a metal stack of 20/100/10 nm of Ti/Au/Ti using an electron-beam assisted evaporation machine (Balzer PLS 570, Pfeiffer, Germany). Afterward, a lift-off process was conducted in an acetone bath for 2.5 h to wash off the sacrificial material and photoresist. Then, a second flexible PaC passivation layer with a thickness of 5 µm was deposited as described before. Next, the flexible polymer was removed from the 3D cut-outs of the *kirigami* design, as well as the outline of the shape of the device, the contact pads, and electrode openings. Here, an etch mask was patterned on top of the last PaC layer by spin-coating the photoresist AZ 12XT (MicroChemicals GmbH, Germany) at 1000 rpm for 180 s with a ramp of 200 rpm/s, performing a soft bake using a hot plate at 110°C for 4 min, and exposing it to a dose of 350 mJ/cm2, a defoc of 2, and a CDB of -800 with UV at 375 nm with a maskless aligner. Afterward, the wafer was subjected to a post-exposure bake using a hot plate at 90°C for 1 min, followed by a developing step of 2 min using AZ 326 MIF. After patterning the etch mask, a reactive-ion etching (RIE) step using an O2/CF4 gas mixture was conducted to etch PaC. A second RIE step was performed to etch the top 10 nm Ti-layer using an O2/Ar gas mixture. After RIE, the etch mask was stripped using AZ 100 remover (MicroChemicals GmbH, Germany) and then rinsed in isopropanol.

The 2D *kirigami* templates were then released from the host silicon wafer using droplets of water and tweezers. The 2D *kirigami* device was then flip-chip bonded on top of a customized printed circuit board (PCB). First, the PCB was pre-heated at 180°C on a direct contact hot plate and the low-temperature solder alloy Sn42/Bi58 (AMTECH, USA) was applied on the contact pads of the board, which allowed the formation of liquid bumps on each contact pad. Lowering the temperature to 160°C, the flexible probes were aligned and placed on top of the liquid solder paste bumps, which solidified after quickly removing the new chip from the hot plate and cooling it down at room temperature. The recently soldered area was sealed with a polydimethylsiloxane (PDMS) coating with a mix ratio of 1:10 and cured at 120°C for 30 min in an oven.

An electrode coating to improve the electrochemical properties of the Au-based electrodes was needed. So, a conductive polymer, PEDOT:PSS (poly (3,4-ethylenedioxythiophene: poly(4-styrenesulfonate)) was electrodeposited. Here, an EDOT:PSS solution was prepared with 3,4-Ethylenedioxythiophene (EDOT) and poly (sodium 4-styrenesulfonate) (PSS) with a 0.1% (w/v) and 0.7% (w/v) concentrations in deionized water. Before the deposition, the probes were first subjected to electrochemical cleaning in 1× PBS (phosphate buffered solution) at room temperature by applying 10 cyclic voltammetry cycles to all electrodes using a sweep rate of 100 mV/s and potential limits between -0.6 – 0.9 V versus a Ag/AgCl reference electrode. Then, the surface of the 3D *kirigami* device was activated with O_2_-plasma using a pressure of 0.8 mbar and a power of 80 W for 3 min. The electrochemical polymerization of the EDOT:PSS on the Au-based electrodes was then performed *via* chronoamperometry using a constant potential of 1V for 20 s.

#### Fabrication of the molds

For the fabrication of the molds, the two-photon polymerization 3D printer Photonic Professional GT2 from NanoScribe GmbH was used. In this process, an erbium-doped femtosecond laser source (center wavelength 780 nm) was focused into a liquid droplet of a photoresin. If the laser power exceeds a certain threshold, the photoresin will polymerize only in the focal spot of the laser and thus really complex structures with high resolution can be achieved. The molds were designed using a CAD software and converted to print job instructions using Describe (Software by NanoScribe GmbH). A Zeiss 25X NA0.8 objective was used and IP-S (NanoScribe GmbH, Germany) as photopolymer material. Using the given print recipe from NanoScribe designed for the combination of the 25X objective and IP-S, the slicing distance was set to 1 µm and the hatching distance was set to 500 nm. A scan speed of 100000 µm/s, a Laser Power of 100 %, and a Power Scaling of 1.2 lead to the best printing result with sufficient resolution and stability of the printed structure. The molds were printed onto commercial 2.5 cm x 2.5 cm ITO-coated glass substrates (Nanoscribe GmbH, Germany) which were previously coated with 3 µm PaC which ensured high adhesion of the print to the glass substrate. A developing step after the printing process washed away the remaining photopolymer which was not polymerized. The samples were placed into a bath of fresh mr-Dev 600 developer for 15 min followed by another 5 min bath in refreshed mr-Dev 600. Finally, the molds were placed into IPA for another 5 min and then air dried.

#### Thermoforming

After the folding of the 2D probe inside the molds, a thermoforming step was carried out to keep the 3D structure after separating the flexible probe from the molds. Researchers propose to conduct thermoforming in a vacuum oven to prevent thermal oxidative degradation^39^. Additionally, a slow temperature ramp is required in the heating and cooling step to prevent failure of the thermoplastic material due to thermal and mechanical stresses. When optimizing a thermoforming protocol, temperature is a more critical factor than time, as it has been observed that the crystallization reaction is completed during the first minutes of the thermoforming process^61^. It is important for the thermoplastic material to have a melting point lower than the glass transition temperature of the material of the mold. Failure can occur due to sticking of the PaC to the molds in case of higher temperatures (> 170°C). On the other hand, if the temperature is too low or the process too short, the shanks fold back down again. Therefore, it is required to find optimal parameters. Considering what was previously mentioned, the thermoforming protocol is conducted as followed:

1. The probe and the molds are placed into the oven at room temperature.
2. Temperature ramp of 5°C/min until final temperature of 160 °C is reached.
3. The maximum temperature of 160°C is held for 60 min.
4. The oven is turned of. Probe and molds are cold down for 120 min.
5. The probe and the molds are carefully separated.

### Mechanical and electrochemical characterization

Simulations of the folding process were done using COMSOL Multiphysics and the shell module. The Young’s modulus of PaC was set to 1.7 GPa, which was determined experimentally with tensile tests of PaC stripes (10 µm thickness, 2 mm widths) using the UniVert tensile tester (CellScale). For the calculation of the critical buckling load Euler’s formula was used.

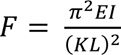

Here, the Young’s modulus (*E*) of PaC was set again to 1.7 GPa, the second moment of inertia is *I*, the column effective length factor *K* = 0.7 for a fixed-pinned boundary condition, and *L* is the length of the shank.

The critical length for self-buckling *L_b_* was calculated following Greenhill’s equation:

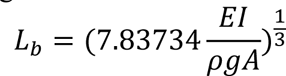

where *E* corresponds to the Young’s modulus, *I* is the second momentum of inertia, *ρ* is the density, *g* the acceleration due to gravity, and *A* corresponds to the cross-section. The calculations were carried out for the protruding blocks of the lower molds as well as for a single PaC-shank. In both cases, a clamped constraint for the base and a free situation at the top of the block or the shank were assumed for the boundary conditions^62^. The material properties of IP-S were given by the manufacturer (*E* = 2.1 GPa, *ρ* = 1.19 g cm^-3^ for solid structures at 20 °C). For the material properties of PaC a Young’s modulus of *E* = 1.70 GPa (Table 1) and a density of *ρ* = 1.289 g cm^-3^ were used^63^. The cross-section *A* was calculated at the bottom of a protruding block in the different designs used. For a rectangular block the momentum of inertia *I* was determined as follows:

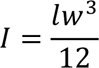

where *l* is the length of the protruding block and *w* is the corresponding width.

Electrochemical impedance spectroscopy (EIS) was measured in 0.1M phosphate-buffered saline (PBS) using a Biologic VSP-300 (Bio-Logic SAS, Claix, France) potentiostat and a three-electrode configuration. Here, each electrode of the 3D *kirigami* MEA was a working electrode, and a Pt wire and an Ag/AgCl pellet were used as counter and reference electrodes, respectively.

### Imaging

Pictures of 2D *kirigami* template before the folding process were taken with a Nikon L200N microscope. Pictures of the 3D printed structures were then taken using a scanning electron microscope (SEM, Gemini 1550 instrument (Leo/Zeiss)). SEM imaging was improved by enhancing the conductivity of the 3D probes by sputtering a thin layer of iridium oxide (6 nm) onto the sample (current 15 mA for 1 min). The imaging was taken at a 3 kV acceleration voltage.

### Tissue preparation and surgery for in vitro and in vivo recordings

#### In vitro testing of human brain slices

Human neocortical brain slice cultures were prepared from access tissue (cortical tissue outside the epileptic focus, resected to gain access to the pathology) obtained from patients undergoing epilepsy surgery. Approval (EK067/20) of the ethics committee of the University of Aachen as well as written informed consent was obtained from all patients. The tissue preparation was performed by the Clinic for Neurology, Uniklinik RWTH Aachen, according to published protocols^64^. To ensure tissue integrity the cortex was carefully micro-dissected and resected with only minimal use of bipolar forceps. The section is cut perpendicular to the surface; thus it contains all six layers of the cortex. It was then directly transferred into ice-cold artificial cerebrospinal fluid (aCSF) (in mM: 110 choline chloride, 26 NaHCO3, 10 D-glucose, 11.6 Na-ascorbate, 7 MgCl2, 3.1 Na-pyruvate, 2.5 KCl, 1.25 NaH2PO4, und 0.5 CaCl2) equilibrated with carbogen (95% O2, 5% CO2) and immediately transported to the laboratory. During the whole process, the tissue was kept submerged in cool and carbogenated aCSF at all times. After removal of the pia, tissue chunks were trimmed perpendicular to the cortical surface and 250– 350 µm thick acute slices were prepared using a Microm HM 650V vibratome (Thermo Fisher Scientific Inc). Afterwards, the slices were cultured according to pre-published protocols^64^ or measured as acute slices. The slices were then transported to the laboratories at IBI-3, Forschungszentrum Jülich in O_2_ saturated aCSF. Once the slices arrived, they were placed into aCSF (in mM) 124 NaCl, 24 NaHC03, 3 KCl, 1.25 NaH2P04, 1.25 MgCl2, 2 CaCI_2_, and 10 glucoses oxygenated with carbogen and placed on a hot plate. For electrophysiological recordings, the brain slices were placed on donut-shaped filter papers, fixed with insect pins, and placed into a perfusion chamber at room temperature.

#### In vivo animal testing

The mice for *in vivo* experiments were bred at the Institute of Biology 2 in Aachen. The experiments in this work have been approved by the *Landesumweltamt für Natur, Umwelt und Verbraucherschutz Nordrhein-Westfalen*, Recklinghausen, Germany, according to German animal protection law and ethics and is reported according to the ARRIVE guidelines.

The mouse strains were obtained from the Jackson Laboratory and subsequently maintained at the local breeding facility at RWTH Aachen University. To facilitate the imaging of calcium-related fluorescence, mice were generated through a double transgenic process involving the crossing of the TRE-GCaMP6s (G6s2, JAX 024742) and Camk2α-tTA (CaMKII-tTA, JAX 007004) mouse strains. This process yielded reliable and consistent expression of the calcium indicator GCaMP6s across all excitatory pyramidal neurons in the cortex of adult mice. Mice were adult males between 12-20 weeks of age and we used two mice for acute and chronic trials, respectively.

### Surgical procedure

During surgery, the mice were placed in a stereotactic frame with monitored body temperature (37 °C) and anesthetized using 1–5% of isoflurane. After an initial cut, the skin at the skull was gently pushed to the side. For low-noise signal recordings, a reference pin was placed on the cerebellum. Then, a craniotomy of about 4 mm in diameter was performed at the left or right hemisphere using a biopsy punch to mark the location and an orthopaedic drill (Eickemeyer) to open the skull. The exposed dura was covered with PBS and cleaned. After an initial incision at the edge of the craniotomy window, a duratomy was performed with a hooked needle to gain access to the cortex. Subsequently, the *kirigami* probes were carefully placed on top of the desired position of the dried cortex using a 3-axis micromanipulator (MTM-3, World Precision Instruments). A metal rod, fixed on a second micromanipulator (uMp4, Sensapex), was used to push the implant to penetrate the tissue. The insertion into the cortex was done with step sizes between 100–250 μm and a velocity of 200 μm s−1. For experiments in the visual cortex, after the successful insertion of a kirigami probe, a glass window was placed on the implant. Subsequently, the implant was fixed on the skull using dental cement. After letting the animal recover, visual stimulation experiments were carried out.

### Widefield imaging

Widefield imaging was done through the cleared intact skull, using a custom-built setup with a tandem-lens macroscope and two 85 mm objectives (Walimex Pro 85mm f/1.4 IF; Walimex). Images were acquired with a sCMOS camera (Edge 4.2, PCO), using a python-based software package (Labcams, https://github.com/jcouto/labcams, by Joao Couto) at a framerate of 30 Hz and a resolution of 512 x 512 pixels. Frames were acquired under alternating illumination using a blue LED (470 nm, M470L3, Thorlabs) and a violet LED (405 nm, M405L3, Thorlabs) and GCaMP6s fluorescence signals were isolated using a 525 nm emission filter (86-963, Edmund optics) in front of the camera. Images acquired under violet illumination captured calcium-independent fluorescence at the isobestic point of GCaMP^65^. Therefore, by subtracting the linearly-rescaled calcium-independent signal from the calcium-dependent signal acquired under blue illumination we could remove intrinsic signal due to hemodynamic fluctuations. The images were then further processed using self-written MATLAB scripts (MATLAB R2019b, MathWorks).

### Sensory stimulation

Tactile stimulation to record neural responses from the somatosensory cortex during either widefield or electrophysiological recordings was performed by a custom-built stimulator, comprised of a rod attached to a small servomotor (TGY-306G-HV) and controlled by a Teensy 3.6 microcontroller (PJRC LLC). By moving the motor at high speeds, the rod applied mild pressure to the hindlimb for a duration of 0.1 seconds every five seconds and stimulation was repeated for a total of 50 trials. To compute cortical responses to tactile stimulation during widefield imaging, we then computed the averaged cortical activity in the first 150 ms relative to tactile stimulation.

To identify the location of visual cortical regions, we performed retinotopic mapping^66^ during widefield imaging. We presented continuous periodically drifting bars with a flickering checker pattern as visual stimuli. Stimuli were created using Psychtoolbox^67^ in MATLAB and composed of a static checker pattern with a patch size of 20° and contrast reversal at 3Hz. This pattern was masked and only visible through a 15°-wide bar aperture, moving over the screen with a temporal period of 0.5 Hz in all 4 cardinal directions in randomized order. Stimuli covered roughly 120° horizontally and 80° vertically of the visual field of the mouse and were pre-rendered in advance. A spherical correction was applied to compensate for distortions due to the presentation on a flat monitor. The monitor (BenQ XL2420T) was positioned at an angle of 20° to the midline and tilted 10° above the animal. The eye was positioned at the horizontal center of the monitor, 30 mm above the lower edge of the monitor and the stimulus presentation was adjusted accordingly. By combining functional responses and anatomical landmarks we could then register the imaging data to the Allen Common Coordinate Framework^51^ to reliably identify the location of the primary visual cortex and the surrounding higher visual areas^68^.

To trigger different neural responses across cortical regions we either presented a simple moving grating stimulus or broadband random phase textures to create more complex visual stimuli^69^. The moving grating stimulus consisted of a full-field horizontal grating with a spatial frequency of 0.04 cpd, moving with a temporal frequency of 1 Hz. The broadband stimulus consisted of a full-field dense mixture of localized moving gratings with random positions, with a range of orientations between 0 and 45 degrees and spatial frequencies between 0.004 and 0.4 cpd. All stimuli were pre-rendered in Matlab and presented for 5 s, followed by a 5 s inter-stimulus interval during which a mean gray screen was shown. Gratings and broadband stimuli were shown in randomized order for a total of 50 repetitions, respectively.

### Sectioning of the brain

Following perfusion of the mouse with PFA (4%), implants were retracted, and the brain was extracted from the skull. The brain was fixed for one day and then prepared for cryo-sectioning. To provide cryoprotection, the brains were immersed in 10% sucrose in 1× PBS overnight and then in 30% sucrose in 1× PBS until sectioning. For sectioning, the frontal lobe and cerebellum were removed, and the brain was embedded in Tissue-Tek® O.C.T.™ compound and frozen. The frozen brain was cut horizontally at 20 µm thickness using a LEICA CM3050 S at -23°C chamber temperature and -21°C object temperature. The slices were collected on Epredia™ SuperFrost Plus™ adhesion slides, dried, and then stored at -80°C until staining.

#### Histology

Rodent as well as human brain slices were stained using the antibodies NeuN (MAB377, MilliporeSigma (Merck), 1:1000; anti mouse A48287 (AF555) 1:750), GFAP (ab5541, MilliporeSigma (Merck), 1:1000; anti chicken A362933 (AF647)), and Iba-1 (19-19741, Wako Chemicals, 1:1000; anti rabbit A32732 (AF488) 1:750), as well as DAPI (D9542-1 mg, MilliporeSigma (Merck) 285, 5 nM). The human brain slices were washed 3 times in PBS, kept for 1 h in 15 % succrose, washed again 3 times with PBS, and then blocked for 2 h with 10 % goat-serum (with PBS-T-T). The samples were then incubated with the first antibodies for three days, then washed again in PBS and after that incubated overnight with the second antibodies. After doing another washing step, the slices were each stained with 250 µl DAPI and then washed again before they were immersed in fluoromount. After defrosting and drying the rodent brain slices, they were processed in a similar way: 3 x washing in PBS, 30 min in PBS-T-T, 3 x washing in PBS, blocking in 10 % goat-serum for 1 h, incubating with the first antibodies overnight, washed, and incubating with the second antibodies for 90 min and washed again. They were then stained with DAPI, washed and dipped in 200 µl TrueBlack, Cat:23007, Biotium (1:20 in 70% Ethanol) for 30 s. After a last washing step, they were immersed in fluoromount.

#### Data processing of the stained images

Images were taken with a confocal laser scan microscope (Zeiss LSM 710) and processed using ImageJ^70^. For the mice brain slices, ROIs were selected using the wand tool. Then, the image was split into channels and the threshold adjusted. The watershed algorithm was used to separate overlaying cells. The particles were counted for the blue (DAPI) and red (NeuN) channel. For the fluorescence intensity the mean grey values were computed inside the ROI as well as inside a reference ROI (500 x 500 µm, placed 500 µm away from implantation site) for the relative fluorescence intensity.

### Electrophysiological data acquisition and signal processing

For *in vitro* as well as for *in vivo* tissue recordings, the ME2100-System (Multi Channel Systems MCS GmbH, Germany) and a 32 channel headstage (ME2100-HS32-M-3m) were used. NeuroNexus adapter (ADPT-NN-16/32 and ADPT-NN-32) served to connect the headstage to our customized 16 or 32 channel PCBs. The McsMatlabDataTools Matlab toolbox ^71^ and self-written scripts were used to import and process offline HDF5 files created by the ME2100-System. The raw traces were bandpass filtered with cut-off frequencies of 100 Hz and 3 kHz using a 6^th^ order zero-phased Butterworth filter to extract spiking activity. For local field potentials (LFPs), the data was low pass filter at 100 Hz. HFOs were filtered using a 10^th^ order Butterworth bandpass filter with cut-off frequencies of 250 and 350 Hz. For tactile as well as visual stimulation, the LFPs were sorted into trials corresponding to the type of stimuli. Then, the responses were averaged (mean ± standard deviation) to be displayed in Figures 5C and 5G. The SNR was calculated using the maximum and averaged peaks of spiking activity divided by the standard deviation of the noise. In order to calculate the maximum value of the cross-correlation during the SLEs in the respective electrode, the raw data trace during the SLEs of the respective electrode was cross-correlated with all other electrodes, and the resulting data were averaged over all SLEs in the trace. This process was repeated for each electrode in Figures 5D.

### Statistical Analysis

Data is displayed in absolute values or mean ± standard deviation if not described otherwise. *N* number of samples are noted for each data evaluation at the respective figure/ table description. To compare the tensile test results of untreated vs. annealed PaC stripes, we conducted an unpaired two-sample t-test with a 95% confidence interval. After confirming the normality of the data using a Lilliefors test, we compared the data between groups to verify the statistical significance of the observed patterns. A paired t-test with 95 % confidence interval was used to compare firing rates upon SLEs in human brain slices.

## Acknowledgments

The authors thank the Helmholtz Nano Facility (HNF) at *Forschungszentrum* Jülich for facilitating the microfabrication of the devices. The authors also thank K. Parker for assisting the execution of tensile tests. The authors thank B. Breuer for organizational support, E. Brauweiler-Reuters for carrying out SEM, and M. Prömpers, M. Banzet, and R. Stockmann for technical support at the HNF. The authors thank B. Gittel for helping with the histological preparation and staining of human and mouse brain slices. This work was supported by the Helmholtz Association (VH-NG-1611) and the *Deutsche Forschungsgemeinschaft* (DFG, German Research Foundation; GRK2610 (project number 424556709) and GRK2416 (project number 368482240).

## Author contribution

M.J. and V.R.M. conceived the development of the *kirigami* probes and planned the study. M.J. fabricated the devices with the support of J.A.S., S.D., L.K., A.O., and V.R.M. J.A.S. and S.D. fabricated the molds for the *kirigami* folding. M.J. characterized the devices with the support of S.D. and L.K. M.J. carried out FEM simulations with the support of S.D., A.O., and V.R.M. A.H. and H.K. supported the retrieval of human brain slices. M.J. carried out *in vitro* experiments with human brain slices with the support of S.M.P, H.K., and V.R.M. S.M. conducted the *in vivo* experiments in mice with the support of M.J., P.S.G., and V.R.M.P.S.G. prepared the mouse brain slices. H.K. carried out the immunohistochemical staining of human and mouse brain slices. M.J., P.S.G., and S.M.P imaged the neural slices. M.J. collected and processed data and figures with the support of S.M., H.K., A.O., and V.R.M. M.J. and V.R.M. wrote the initial draft of the manuscript. All authors reviewed and edited the manuscript. V.R.M. supervised the project.

