## Supplementary material for "Flexible 3D *kirigami* probes for *in vitro* and *in vivo* neural applications"

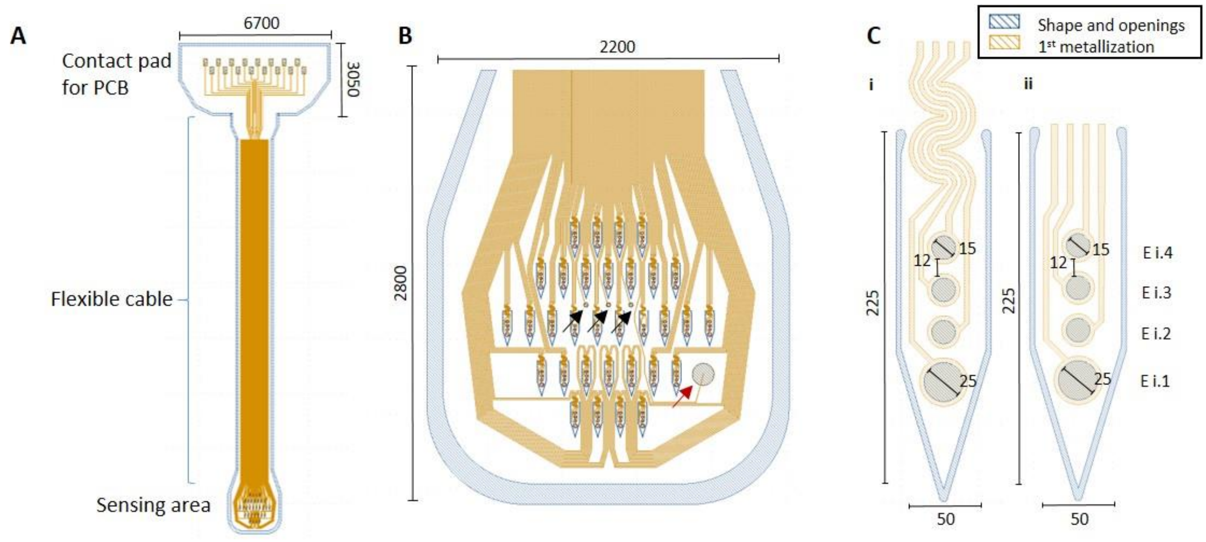


**Figure S1: 3D *kirigami* MEA design.** Exemplary design including a contact pad region, a flexible cable, and a sensing area (A) containing surface electrodes (arrows) and penetrating shanks (B), with either meander (Ci) or straight feedlines (Cii). Surface electrodes in (B) can be either used for recording (black arrows) or as an internal reference electrode (red arrow).

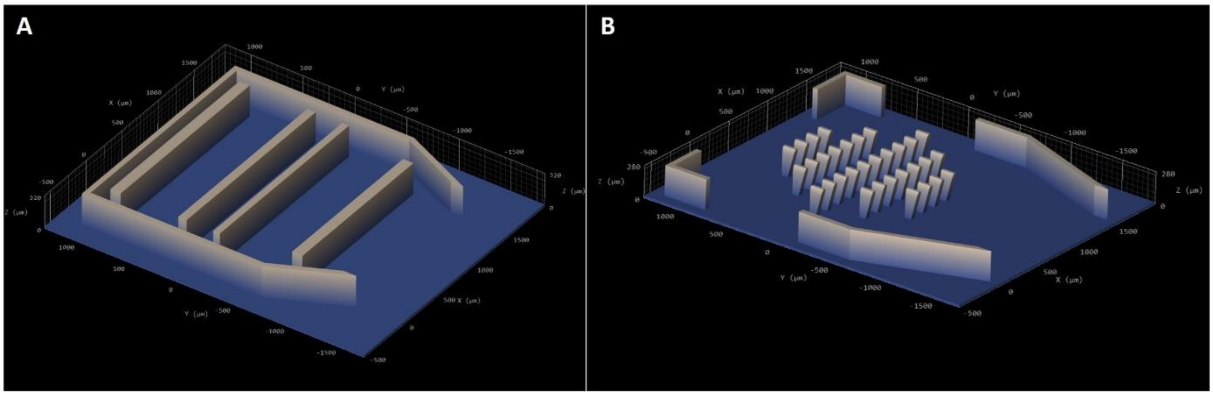


**Figure S2: Molds for *kirigami* matched die forming process.** Exemplary upper (A) and lower (B) molds for folding 2D *kirigami* MEAs with 32 shanks of a length of 225 µm. Lower and upper molds both contain sidewalls that are used as alignment structures upon folding. The upper mold (A) additionally contains elongated protruding structures to ensure homogeneous pressure during folding. The lower mold (B) contains protruding structures (blocks) which match the location of the shanks to press them out of the 2D plane.

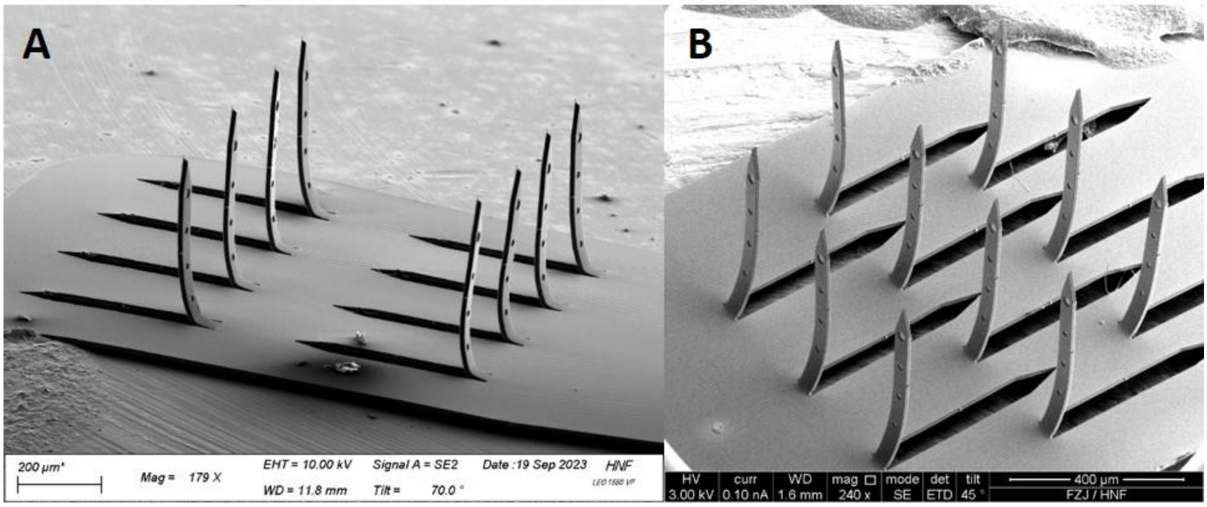


**Figure S3: 3D *kirigami* MEA designs.** Exemplary array designs with parallel shank rows (A) and diamond-shaped design (B).

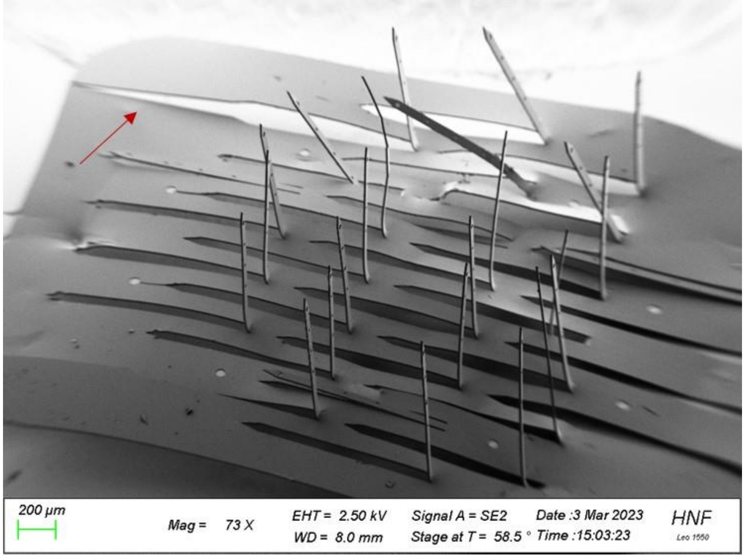


**Figure S4: Mechanical instability of 2D template in high density shank arrays.** Ripping of the 2D layout of a probe with 30 x 1000 µm long shanks and an inter-shank distance of 120 µm.

**Table S1: Mechanical properties of untreated (no annealing), annealed at 160°C, and annealed at 200 °C 10 µm thick PaC stripes.** Values are displayed as mean +/- standard deviation. p-values < 0.05 are marked with * (N = 7 or 8 for each group, used statistical method: one-way ANOVA with 95 % confidence interval).

| | Untreated (no annealing) | Annealed at 160°C | Annealed at 200°C |
|---|---|---|---|
| Young's modulus (GPa) | 1.66 +/- 0.22 | 1.70 +/- 0.32 | 2.08 +/- 0.34* (p = 0.03) |
| Stress at fracture (MPa) | 83.82 +/- 2.02 | 89.07 +/- 6.92* (p = 0.034) | 91.45 +/- 6.3* (p = 0.006) |
| Strain at fracture (%) | 14.15 +/- 4.65 | 13.93 +/- 11.25 | 9.56 +/- 9.5* (p = 0.022) |

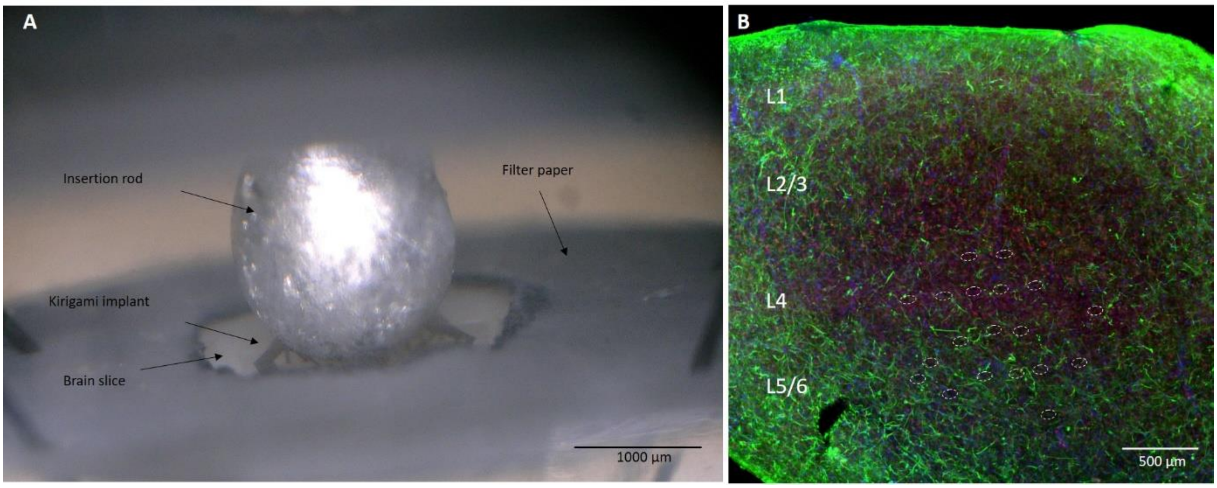


**Figure S5: Insertion of a kirigami implant into a human brain slice.** A) Insertion of a kirigami implant into an *in vitro* human brain slice using an insertion rod. The brain is fixated in a perfusion chamber with the help of filter paper and insect pins and embedded in aCSF. B) Stained brain slice and marked insertion holes (Green: GFAP, blue: DAPI, red: NeuN)

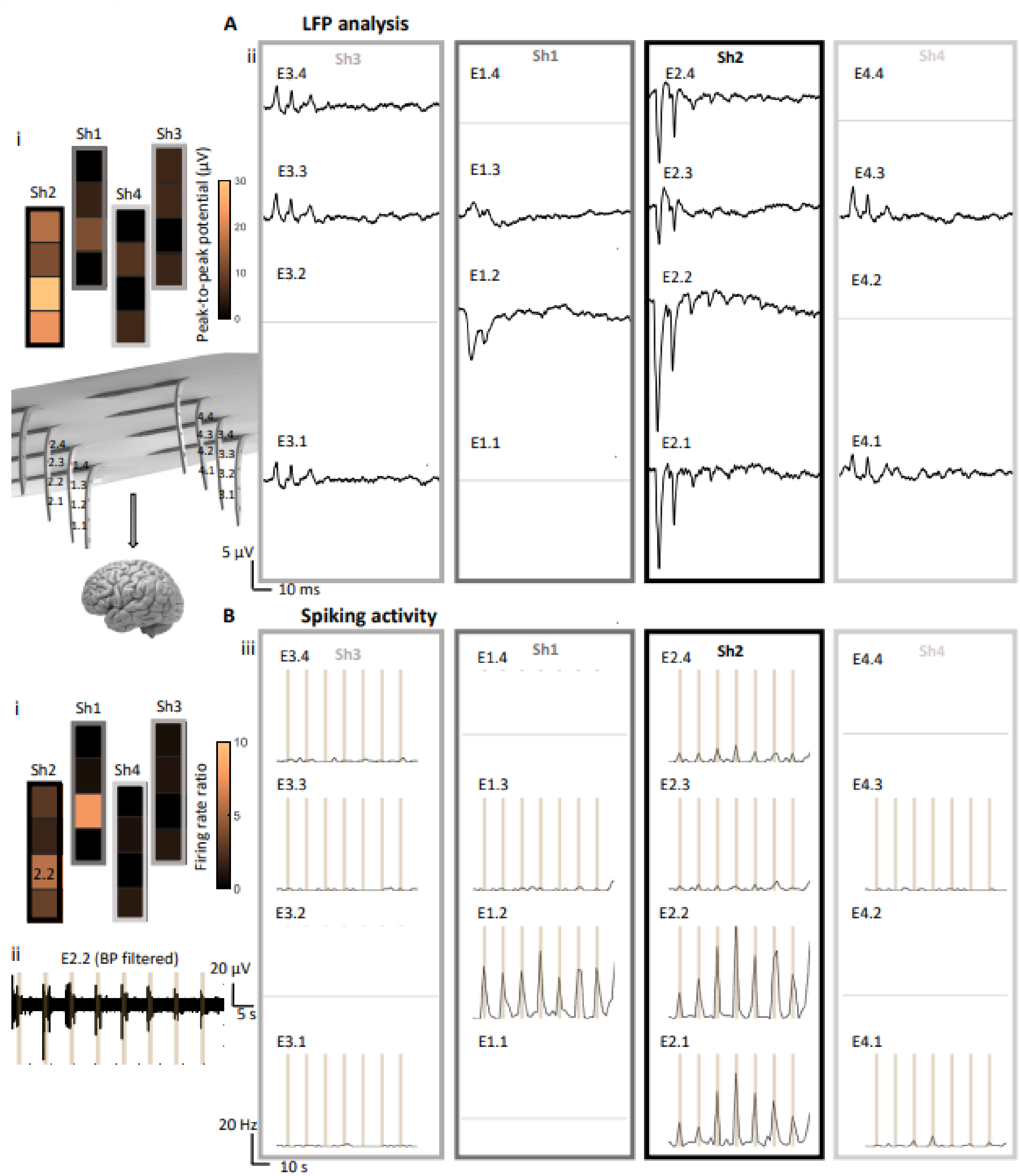


**Figure S6: 3D electrophysiological recordings of acute in vivo mouse somatosensory cortex.** A) Averaged LFPs show the spatial activity spread depending on the depth (z) and x-y location of the electrodes(ii). The peak-to-peak potentials were based on the averaged LFPs over 50 trials (i). B) Average changes in the spiking activity equally differs according to the electrodes position. Shown are changes in firing rates due to tactile stimulation (i), an example of the band pass filtered data of electrode E2.2 (ii) and the firing rate traces for all electrodes (iii). The firing rates rise respectively with the stimulation (brown bars). The broken electrodes are marked with grey lines.

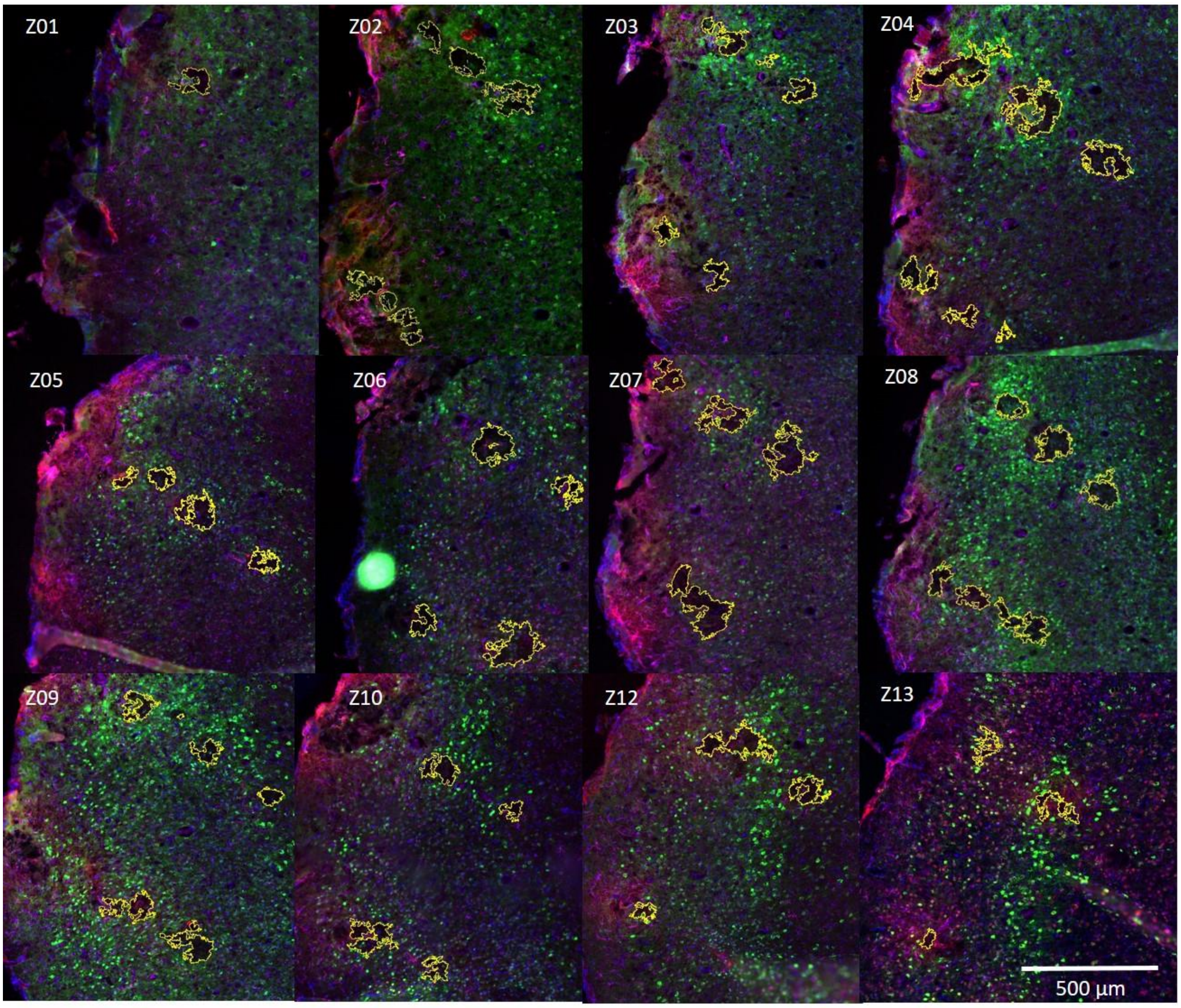


**Figure S7: Immunohistological analysis.** Stainings for mature microglia (IBA1, green), cell nuclei (DAPI, blue), microglia (GFAP, magenta) and mature neurons (NeuN, red) of 20 µm-thick horizontal slices after brain perfusion and retraction of a *Kiri*-500 implant chronically implanted for four weeks in the visual cortex of a rodent. Outlines in yellow denote the regions of interest (ROIs) enclosing the implantation lesion. Concentric or eccentric ROIs indicate a small ROI enclosing the direct implantation footprint of a shank, and a large ROI enclosing visible foreign body reactions (FBRs).

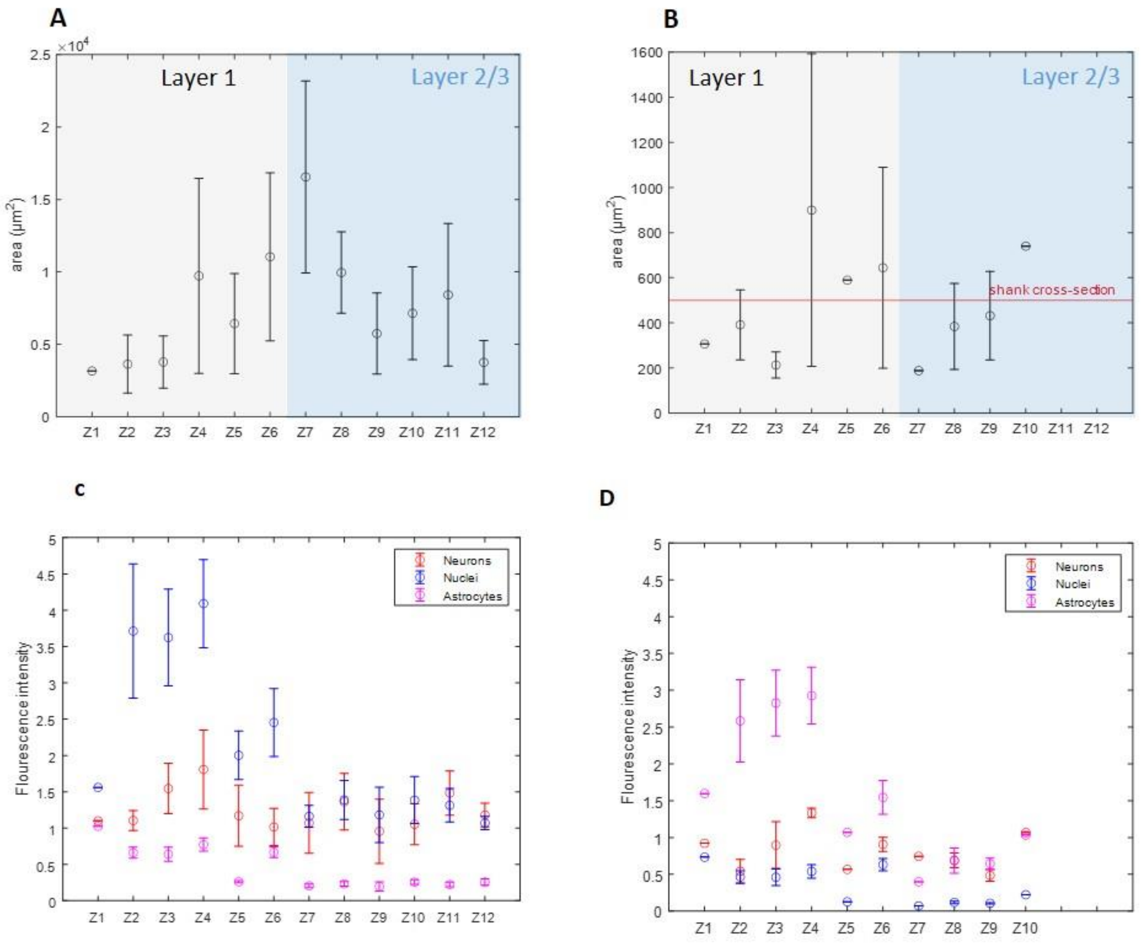


**Figure S8: Sizes of ROIs and relative fluorescence in immunohistological stainings.** The size of the large (A) ROIs depends on the z-depths inside the cortical layer (Z1 closest to surface). In cortical layer I, the affected regions are smaller than in layers II and III. In contrast, the size of the small (B) ROIs (insertion lesion) is smaller than the shank's cross-sections. The relative fluorescence is overall increasing in deeper layers. Microglia in the large (C) and small (D) is getting more and more similar to the reference ROI (rel. intensity ~1), while there is also a growing amount of neurons and nuclei.

**Supplementary Table S2**: **Literature comparison with Utah and Michigan array as well as other kirigami approaches.**

| Publication | Dimension | Electrode count | Shanks |
|---|---|---|---|
| Campbell et al, 1991 (Utah array) | 1500 µm length<br>90 µm diameter at the base | Typically 64, up to 1024 | 64 |
| Wise et al., 2004 (Michigan array) | Typically 25 – 50 µm shank width | Up to 1024 electrodes, 128 Ch | 256 |
| Takeuchi et al., 2003 | 1200 µm long shanks<br>160 µm wide<br>20 µm thick | 18 | 6 |
| Sim et al., 2018 | ~1600 mm long<br>~ 300 µm wide<br>7.65 µm thick | 64 | 16 |
| Chen et al., 2010 | 3500 mm long<br>200 µm wide<br>27.5 µm thick | 8 | 4 |
| Soscia et al., 2020 | 1100 µm long<br>90 µm wide<br>15 µm thick | Chip with up to 256 electrodes containing arrays with up to 80 electrodes | 10 |
| Lee et al., 2022 | 1500 µm length<br>~200 µm wide<br>< 20 µm thick | 24 penetrating electrodes,<br>9 surface electrodes | 4 |
| This work | Up to 1000 µm length<br>50 µm wide<br>10 µm thick | Up to 512 electrodes | Up to 128 |

### Supplementary videos

**SV1** – Folding of a *KiRi*-225 probe

The video shows how a *KiRi*-225 is folded. The flexible probe is placed inside a 3D printed lower mold. When placing the upper mold on top, all shanks fold at the same time.

**SV2 –** Insertion of *Kiri*-500 probe into agarose gel.

The insertion of a *KiRi*-500 probe was tested using agarose gel which mimics the mechanical properties of neural tissue.

**SV3** – Insertion of *Kiri*-500 probe into mouse cortex.

Insertion of a *KiRi*-500 into the mouse cortex was performed after removal of the dura. The 500 µm long shanks are inserted using an insertion rod, enabling a single-shot insertion to reach layer 2/3 of the mouse cortex.
